# Peripheral T Helper Cells Dominate the Synovial CD4^+^ T Cell Compartment in Systemic Juvenile Idiopathic Arthritis and Are Shaped by IL-1β and IL-18

**DOI:** 10.1101/2025.07.03.661735

**Authors:** Johannes Dirks, Jonas Fischer, Leon Harrer, Claudia Bracaglia, Giusi Prencipe, Manuela Pardeo, Silvia Magni-Manzoni, Ivan Caiello, Annette-Holl Wieden, Hermann Girschick, Christoph Kessel, Henner Morbach

## Abstract

**Objectives:** Systemic juvenile idiopathic arthritis (sJIA Still’s disease) is primarily driven by systemic autoinflammation but can progress to chronic arthritis, implicating dysregulated adaptive immunity. However, the molecular mechanisms within inflamed joints that link innate and adaptive immune activation remain poorly defined. This study aimed to define the transcriptional and clonal landscape of synovial CD4^+^ T cells in sJIA and to identify inflammatory signals driving their differentiation.

**Methods:** Synovial fluid CD4^+^ T cells from patients with sJIA and oligo/polyarticular JIA (o/p-JIA) were analyzed using flow cytometry, scRNA and TCR sequencing. Cytokine secretion and B helper function were assessed *in vitro*. The effects of IL-1β and IL-18 on T cell differentiation were evaluated using bulk RNA sequencing and multiplex ELISA.

**Results:** sJIA joints harbored a dominant population of clonally expanded CD4^+^ T cells with a peripheral T helper (Tph) cell state, marked by IL-21 and CXCL13 expression and robust B cell help. scRNA-seq revealed a heterogeneous CD4^+^ T cell landscape, with transcriptional convergence of Tph and regulatory (Treg) programs particularly in sJIA. A subset of these cells exhibited molecular features consistent with differentiation from Tph precursors. *In vitro*, IL-1β and IL-18 promoted Tph differentiation, aligning with the transcriptional profiles of expanded effector Tph cells observed in sJIA joints.

**Conclusion:** These findings identify Tph cell-driven adaptive immunity as a key feature of chronic arthritis in sJIA and link IL-1β and IL-18 to Tph cell induction. This challenges the classical autoinflammatory paradigm and suggests that early cytokine-targeted therapy may modulate T cell fate and disease course.

## Introduction

Systemic juvenile idiopathic arthritis (sJIA, Still’s disease) is a distinct subtype of childhood arthritis characterized by pronounced systemic inflammation, sharing clinical and immunological features with autoinflammatory syndromes. The initial presentation typically includes fevers and rash. Notably, arthritis may be absent at disease onset but frequently develops over time, potentially progressing into chronic, destructive joint disease similar to other subtypes of JIA that are considered autoimmune-driven (1, 2).

sJIA is suggested to follow a biphasic model, beginning with an innate immunity–driven systemic phase that may evolve into a chronic phase characterized by dysregulated activation of the adaptive immune system (3, 4). While this paradigm is increasingly supported by clinical and immunogenetic evidence, the molecular mechanisms guiding the transition from innate to adaptive immunity - particularly within the inflamed joint microenvironment - remain poorly understood. IL-1β and IL-18 have emerged as central mediators of systemic inflammation in sJIA, and IL-1-targeted therapies have demonstrated particular efficacy during early stage (5–9). In line with the proposed biphasic model and the involvement of the adaptive immune system in the later phase, the association with HLA class 2 alleles (e.g. HLA-DRB1*11) implicates T helper cells in the pathogenesis of sJIA (10, 11).

IL-1β is known to promote Th17 differentiation from naïve CD4^+^ T cells, and elevated levels of IL-17, as well as increased frequencies of IL-17-producing γδ T cells and CD4^+^ T cells, have been observed in peripheral blood of patients with acute sJIA (12–17). Initially restricted to regulatory T cells, Th17 polarization in sJIA has been suggested to extend to effector T cells over time, culminating in a Th17-skewed effector phenotype (14). Beyond Th17 polarization, naïve CD4^+^ T cells from sJIA patients exhibit a propensity to differentiate into IL-21–secreting cells *in vitro*, phenotypically and functionally resembling peripheral T helper (Tph) cells (18). Tph cells are increasingly recognized as key drivers of local inflammation and B cell help in autoimmune diseases (19). Notably, expansion of Tph cells has been demonstrated in the joints of patients with oligoarticular and polyarticular JIA (o/p-JIA), particularly among those with antinuclear antibody (ANA) positivity (20, 21).

Despite accumulating indirect evidence implicating CD4^+^ T helper cell subsets in the chronic phase of sJIA, most insights to date have been derived from studies of peripheral blood. As a result, the specific identities, transcriptional programs, and clonal dynamics of CD4^+^ T cells within inflamed joints remain largely uncharacterized. This study aimed to define the transcriptional landscape and clonal architecture of synovial fluid (SF) CD4^+^ T helper cells in sJIA and to compare these molecular features with those observed in ANA+ o/p-JIA, which represent prototypical autoimmune phenotypes.

## Methods

JIA patients with active disease undergoing joint puncture for intraarticular steroid injection were included in this study (Supplementary Table 1). Mononuclear cells were isolated from SF and stored in liquid nitrogen until use. Experimental procedures are described in the Supplementary Methods.

## Results

### Activated PD-1^+^HLA-DR^+^ CD4^+^ T cells are expanded in the joints of sJIA patients and exhibit a Tph-like phenotype

To investigate the functional state of CD4^+^ T cells in the inflamed joints of sJIA, we first assessed their expression of PD-1 and HLA-DR - markers indicative of recent activation. PD-1^+^HLA-DR^+^ CD4^+^ T cells were significantly enriched in the SF of o/p-JIA patients compared to those with enthesitis-related arthritis (ERA-JIA), suggesting that local CD4^+^ T cell activation is not merely a byproduct of joint inflammation, but rather a characteristic feature of specific JIA subtypes (Figure 1A, B). Strikingly, their frequencies were even higher in sJIA, exceeding those observed in o/p-JIA (Figure 1B). In the limited number of patients for whom early disease SF samples were available, these activated T cells were already present at high frequencies - supporting the notion that their expansion is an early event in disease pathogenesis, rather than accumulation in the joints over time (Figure 1B).

**Figure 1 –.**
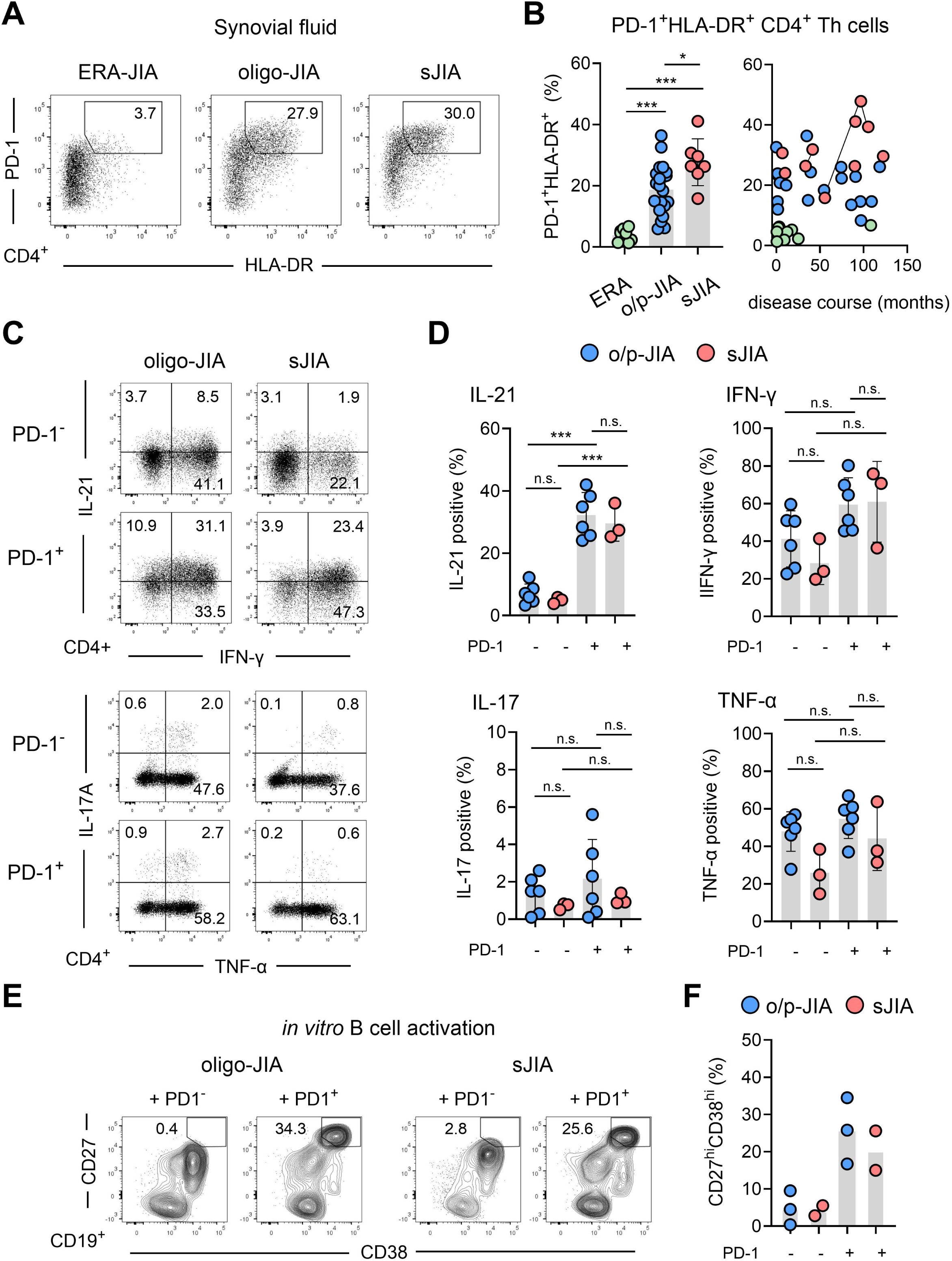
Expansion of IL-21^+^ producing PD-1^+^HLA-DR^+^ CD4+ T helper cells in the synovial fluid of sJIA patients. (A) Representative flow cytometry dot plots showing PD-1 and HLA-DR co-expression on synovial fluid (SF) CD4+ T cells from patients with enthesitis-related arthritis (ERA-JIA), oligo-/polyarticular JIA (o/p-JIA), and systemic JIA (sJIA). (B) Quantification of SF PD-1^+^HLA-DR^+^ CD4^+^ T cells by disease subgroup (left) and correlation with disease duration (right). Disease onset is defined as start of arthritis in ERA-JIA and o/p-JIA and start of systemic inflammation in sJIA. (C) Representative intracellular cytokine staining for IL-21, IFN-γ, IL-17, and TNF-α in PD-1^+^ vs. PD-1^−^ CD4^+^ T cells from o/p-JIA and sJIA SF. (D) Quantification of cytokine-producing cells in SF PD-1^+^ and PD-1^−^ subsets. (E) Representative plots of CD27^hi^CD38^hi^ plasma cells derived from healthy donor B cells after co-culture with SF PD-1^+^ or PD-1^−^ CD4^+^ T cells from o/p-JIA or sJIA patients. (F) Quantification of plasma cell differentiation induced by SF PD-1^+^ and PD-1^−^ CD4+ T cells *in vitro*. Comparisons between groups, as shown in the figure, were performed using Ordinary one-way ANOVA with Bonferroni correction for multiple comparisons. Statistical significance is indicated as follows: p < 0.05 (*), p < 0.01 (**), p < 0.001 (***); n.s., not significant.

Given prior associations of PD-1^+^HLA-DR^+^ CD4^+^ T cells with Tph cell phenotypes in autoimmune contexts (21, 22), we next characterized their cytokine expression profile. IL-21, a signature Tph cytokine, was predominantly expressed by PD-1^+^ CD4^+^ T cells. Quantitative analysis confirmed that IL-21 expression was significantly enriched in PD-1^+^ compared to PD-1^−^ CD4^+^ T cells in both sJIA and o/p-JIA (Figure 1C, D). In contrast, IFN-γ, IL-17, and TNF-α were not preferentially expressed in PD-1^+^ CD4^+^ T cells, with IL-17^+^ cells being particularly scarce (Figure 1C, D). To evaluate their functional capacity, we tested whether PD-1^+^ CD4^+^ T cells could support B cell differentiation *in vitro*. Indeed, PD-1^+^CD4^+^ T cells purified from both o/p-JIA and sJIA demonstrated a robust ability to induce plasma cell differentiation, underscoring their functional resemblance to Tph cells (Figure 1E, F). Together, these findings identify a population of Tph-like CD4^+^ T cells enriched in the joints of sJIA patients.

### Single-cell transcriptomic profiling reveals Tph-enriched but heterogeneous CD4^+^ T cell states in sJIA

To further characterize the synovial CD4^+^ T cell landscape, we performed single-cell RNA sequencing (scRNA-seq) combined with TCR sequencing on CD4^+^ T cells isolated from SF of sJIA and o/p-JIA patients (Supplementary Table 2). After quality control, we analyzed 19,993 CD4^+^ T cells, including 10,216 from o/p-JIA (n=4) and 9,777 from sJIA (n=4).

Unsupervised UMAP clustering identified 21 transcriptionally distinct CD4^+^ T cell clusters, with similar representation across patients (Figure 2A; Supplementary Figure 1). A prominent cluster (cluster 9) expressed high levels of *PDCD1* (PD-1) and *CXCL13*, characteristic of Tph cells (Figure 2A–C). A second Tph-like cluster (Cluster 2) co-expressed *PDCD1*, *HLA-DRB1*, *KLRB1* (CD161), and *ALOX5AP*, but lacked *CXCL13*. Clusters 0, 11, and 19 displayed strong regulatory T cell (Treg) signatures with high *FOXP3* expression, while clusters 7 and 8 expressed cytotoxic genes such as *GZMA* and *GZMK*. Cluster 3 showed a Th17-like profile, and clusters 4, 5, 6, and 18 co-expressed *IL7R*, *SELL*, *TCF7*, and low levels of *CCR7*, consistent with a central memory or naïve phenotype. Cluster 13 represented a proliferative population marked by *MKI67*. Beyond lineage identity, clusters were also defined by gradations of activation, with increasing “effectorness” toward Tph, cytotoxic, and some Treg clusters (Figure 2D) (23).

**Figure 2 –.**
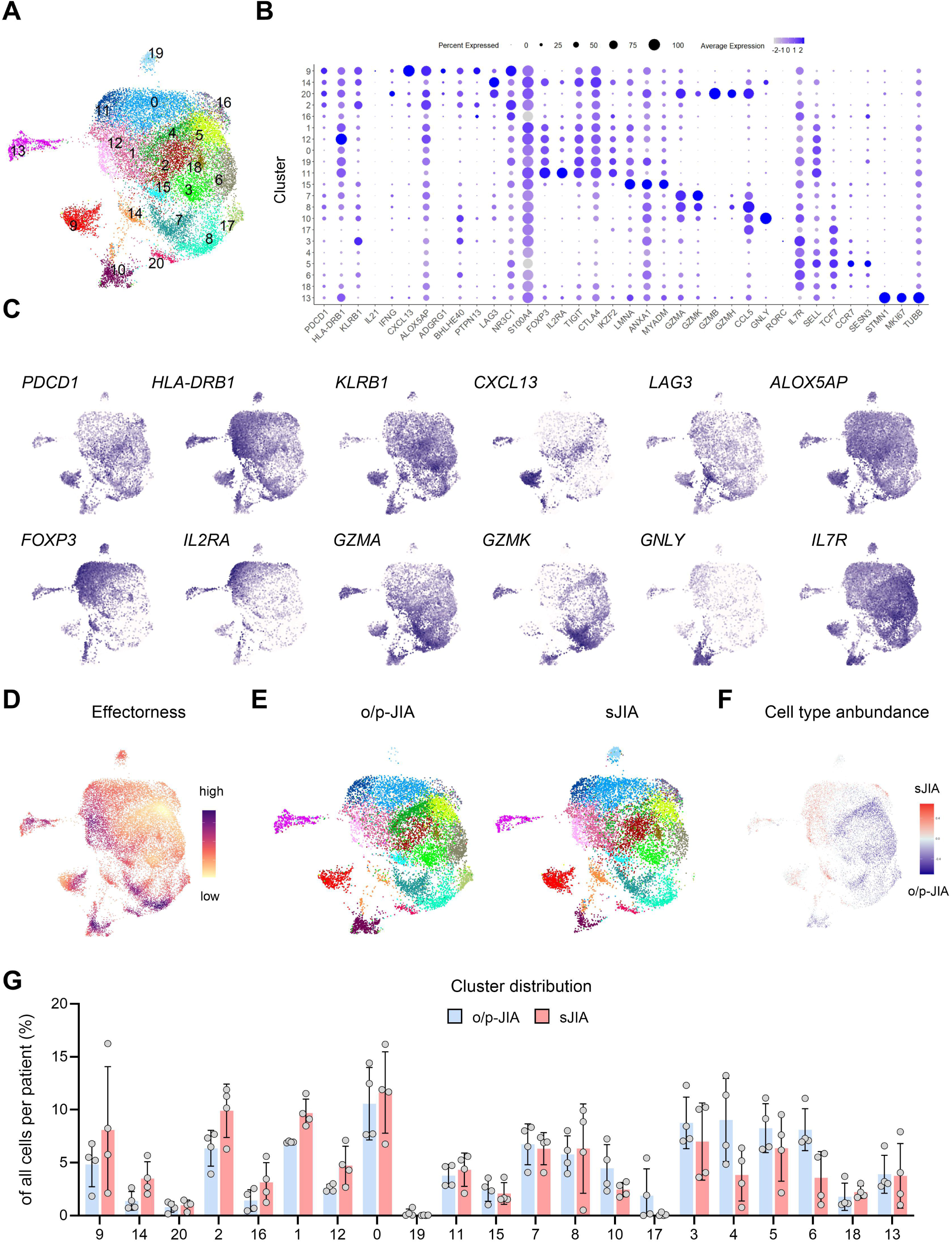
Transcriptional landscape of synovial fluid CD4^+^ T cells in sJIA and o/p-JIA. (A) UMAP projection of synovial fluid CD4+ T cells from sJIA (n=4) and o/p-JIA patients (n=4), with unsupervised cluster annotation. (B) Dot plot showing key cluster-defining and lineage-relevant genes, including transcription factors. (C) Feature plots of genes associated with Tph (*PDCD1, CXCL13, ALOX5AP*), Treg (*FOXP3, IL2RA*), cytotoxicity (*GZMA, GZMK*), and activation (*HLA-DRB1*). (D) UMAP colored by T helper effectorness score. (E) Disease-specific UMAPs showing cluster distribution in sJIA vs. o/p-JIA. (F) UMAP colored by the correlation of covarying neighborhood abundance to sJIA disease status (red = high, blue = low correlation; grey = not significant at FDR<0.05) (G) Quantification and comparison of cluster proportions between sJIA and o/p-JIA. Statistical comparison between groups was performed using multiple unpaired Student’s t-tests with correction for multiple comparisons by Bonferroni-Dunn. No statistically significant differences were found between the groups (all p > 0.05).

Although the overall distribution of CD4^+^ T cell clusters did not differ significantly between sJIA and o/p-JIA, we observed a trend toward increased abundance of Tph-associated clusters in sJIA (Figure 2E-G). Covarying neighborhood analysis (CNA) further identified cell populations enriched in sJIA that spatially mapped to clusters 12 and 14 characterized by transcriptional Tph but also Treg programs, which resist straightforward classification into canonical subsets (Figure 2F). These findings not only corroborate our flow cytometry data, confirming an enrichment of Tph-like CD4^+^ T cells in sJIA SF, but also suggest an expansion of CD4^+^ T cells with mixed transcriptional identities in this disease. The convergence of multiple transcriptional programs likely reflects both the inherent plasticity of CD4^+^ T cells in the inflamed joint microenvironment and the limitations of rigid clustering approaches in scRNA-seq, which may fail to resolve continuous gene expression programs.

### Dominant and overlapping Tph transcriptional programs shape the synovial CD4^+^ T cell landscape in sJIA

To capture the continuum of CD4^+^ T cell states beyond discrete clustering, we applied the T-CellAnnoTator (TCAT) pipeline to our dataset (24). TCAT quantifies pre-defined consensus gene expression programs (cGEPs) - derived via consensus non-negative matrix factorization (cNMF) - that reflect distinct CD4^+^ T cell subsets, activation states, and functional profiles. Unlike hard clustering, cNMF allows individual cells to express multiple gene programs, providing a more granular view of transcriptional heterogeneity (25).

The spatial distribution of Tph-, Treg-, cytotoxic-, and Th17-associated cGEPs across the UMAP paralleled canonical marker gene expression (Figure 2C and 3A). As anticipated, several clusters co-expressed multiple cGEPs, consistent with functional overlap (Supplementary Figure 2). Median expression levels of both Tph and Treg cGEPs were consistently higher in sJIA (Supplementary Figure 3). Notably, cluster 10 - characterized by high *GNLY* expression and CD4^+^ T cell effectorness - exhibited strong Tph cGEP activity uniquely in sJIA, whereas Th17 cGEP expression was significantly elevated in o/p-JIA, highlighting divergent effector polarization across diseases (Figure 3A; Supplementary Figure 3).

**Figure 3 –.**
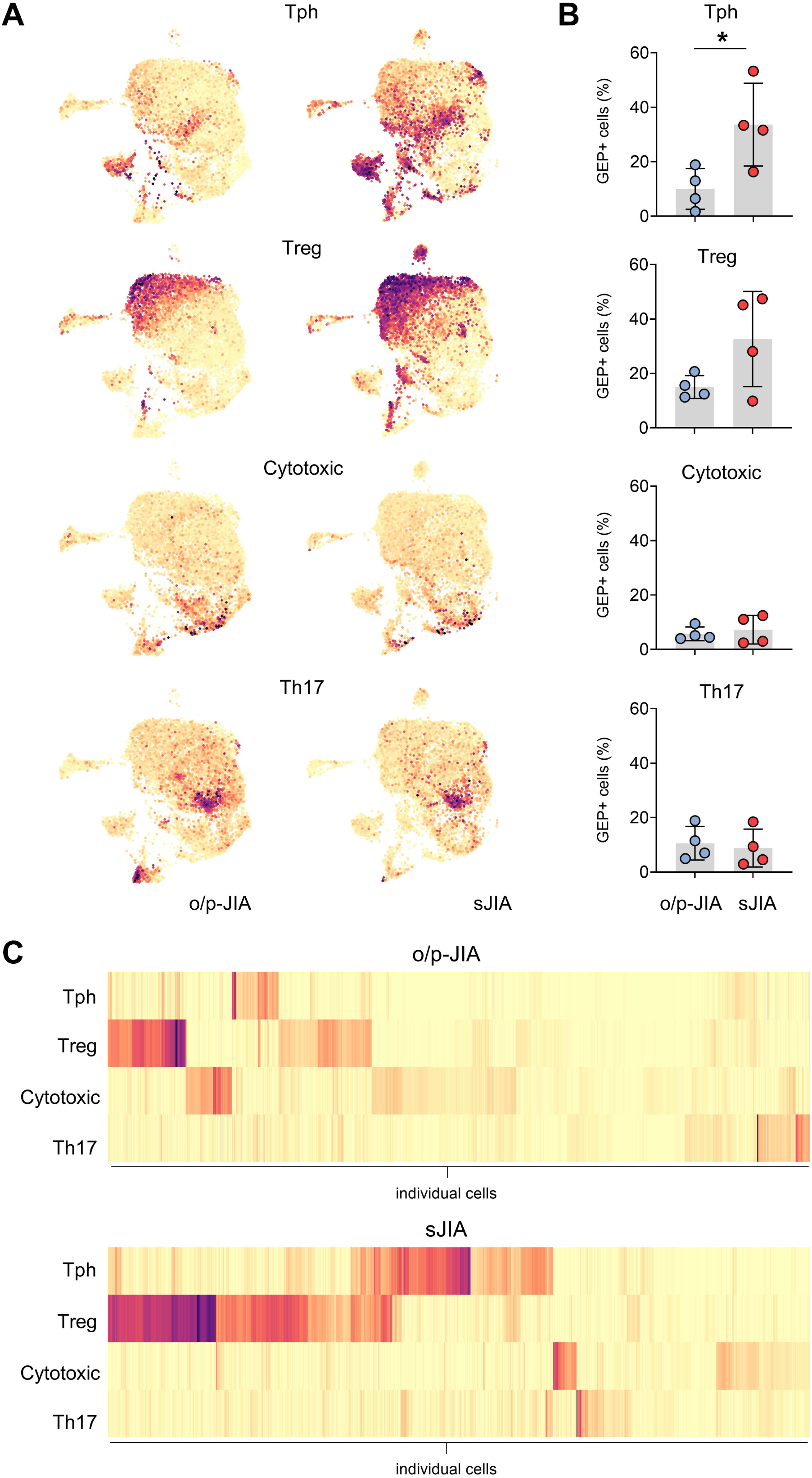
Gene expression programs in synovial fluid CD4^+^ T cells in sJIA reveal a dominant and overlapping Tph signature. (A) UMAPs from Figure 2E, colored by selected consensus gene expression program (cGEP) usage. (B) Proportion of cells with GEPs above defined thresholds. Tph >0.05, Treg >0.1, Th17 >0.05, or cytotoxic >0.05. Statistical comparison between groups was performed using multiple unpaired Student’s t-tests (C) Unsupervised clustering based on per-cell cGEP scores, stratified by disease subgroup using Ward’s minimum variance method with squared Euclidean distances. Columns were divided into 9 clusters.

To enable an unbiased assessment of cGEP activation, we quantified the proportion of CD4^+^ T cells exceeding a defined threshold for each cGEP (Supplementary Figure 3), independent of prior cluster identity. Tph cGEP-positive cells were significantly enriched in sJIA, while Treg cGEP-positive cells showed a non-significant trend toward higher frequency (Figure 3A and B). Th17- and cytotoxic-cGEP frequencies did not differ significantly between diseases (Figure 3A and B).

To further resolve disease-specific transcriptional patterns, we performed unsupervised classification of cells within each disease based on per-cell cGEP expression. In o/p-JIA, Tph- and Treg-like cells segregated into distinct subsets (Figure 3C). In contrast, sJIA harbored a subset co-expressing both Tph and Treg programs, suggesting that CD4^+^ T cells in sJIA not only favor Tph-like states but also exhibit transcriptional convergence of helper and regulatory programs.

### Clonal relationships and phenotypic divergence of CD4^+^ T cells co-expressing Tph and Treg signatures

To dissect the differentiation trajectories of CD4^+^ T cells co-expressing Tph and Treg programs in sJIA, we stratified SF CD4^+^ T cells based on exclusive or overlapping expression of these transcriptional signatures into four groups: Tph-only (Tph^+^Treg^−^), Treg-only (Tph^−^Treg^+^), double-positive (Tph^+^Treg^+^), and double-negative (Tph^−^Treg^−^) cells (Figure 4A, Supplementary Figure 4). Although less abundant than Tph-only cells overall, double-positive cells were significantly more frequent in sJIA than in o/p-JIA, suggesting that they contribute to the expanded Tph cell pool in sJIA joints (Figure 4B). To investigate their clonal origins, we focused on double-positive cells that shared TCR clonotypes with at least one other group to infer lineage relationships. In o/p-JIA, double-positive cells displayed broad clonal overlap with all subsets. Few showed exclusive clonal relationships to Treg-only cells, suggesting a potential Treg-derived origin of the latter (Figure 4C). In contrast, sJIA exhibited a distinct clonal architecture. While fewer double-positive cells showed multi-subset overlap, a slightly higher proportion shared clones exclusively with Treg-only cells, indicating a potential origin from Treg-like precursors. Most remarkably, a subset of double-positive cells in sJIA displayed exclusive clonal relationships with Tph-only cells - suggesting an alternative differentiation pathway linked to Tph cells, a feature not observed in o/p-JIA (Figure 4C).

**Figure 4 –.**
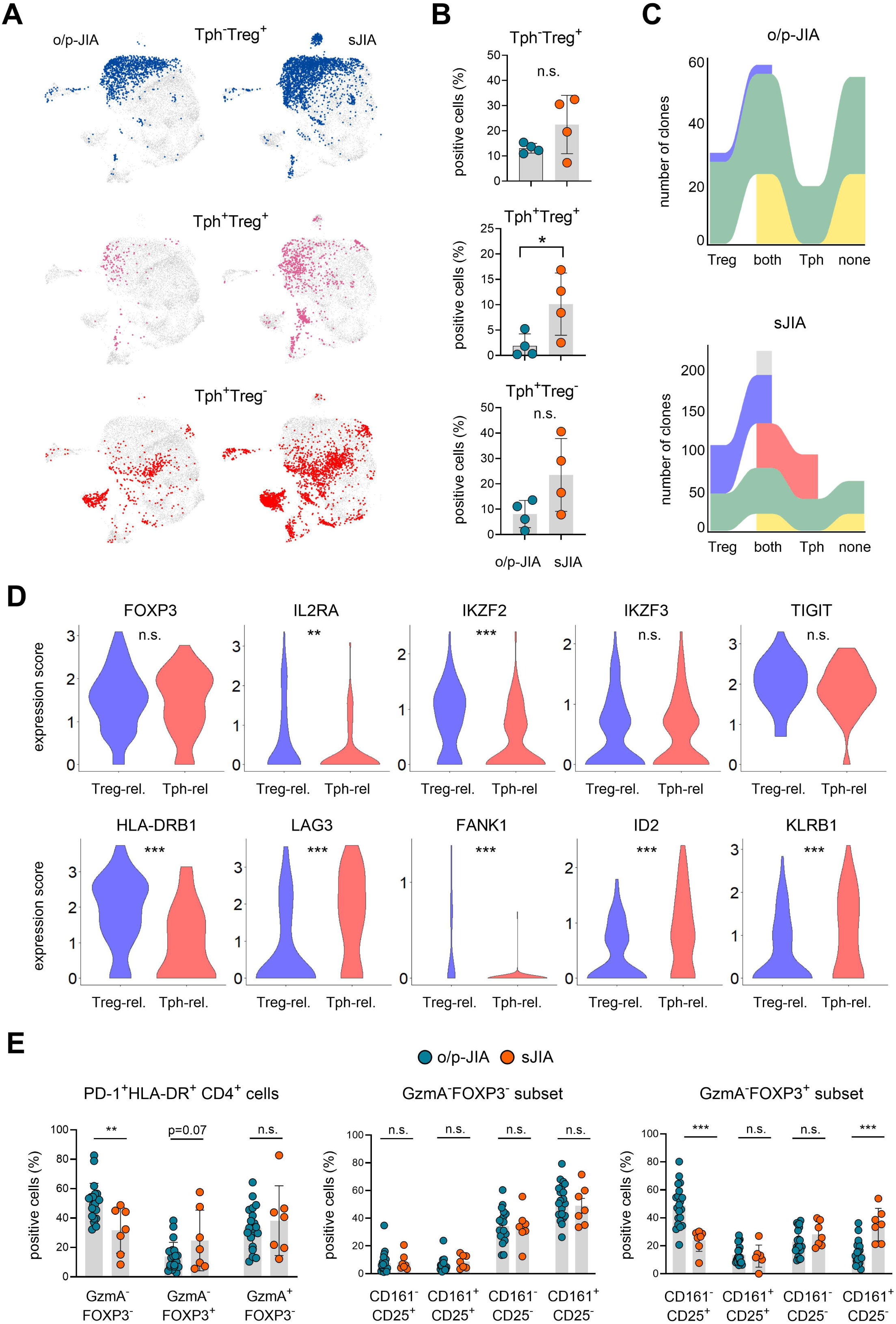
Clonal and phenotypic characterization of CD4^+^ T Cells co-expressing Tph and Treg programs. (A) UMAPs highlighting cells expressing Tph and/or Treg consensus gene expression programs (cGEP) above threshold. (B) Frequency of cells expressing Tph and/or Treg cGEPs. (C) Clonal overlap between cells co-expressing Tph and Treg cGEPs (Tph+Treg+, “double positive”) and other subsets. Only cells that shared clonotypes with at least one other group were included in this analysis. (D) Differential expression of selected genes between Tph^+^Treg^+^ double-positive cells clonally related to either Tph^+^Treg^−^ (“Tph-related”) or Tph^−^Treg^+^ (“Treg-related”) cells in sJIA. (E) Flow cytometric analysis of synovial fluid CD4+ T cells from o/p-JIA and sJIA patients showing frequencies of granzyme A (GzmA) and/or FOXP3 positive subsets among PD-1^+^HLA-DR^+^ cells (left), CD161 and/or CD25 positive cells among the PD1^+^HLA-DR^+^ GzmA^−^FOXP3^−^ (middle) and GzmA^−^FOXP3^+^ T cells (right). Comparisons between groups, as shown in the figure, were performed using two-tailed Student’s t-test in panel B and E. Statistical significance is indicated as follows: p < 0.05 (*), p < 0.01 (**), p < 0.001 (***); n.s., not significant.

To further dissect the double-positive subsets in sJIA, we performed differential gene expression analysis, comparing double-positive cells with exclusive clonal relationship to either Tph-only cells (“Tph-related”) or Treg-only cells (“Treg-related”) (Supplementary Figure 5). While both groups expressed the canonical Treg transcription factor *FOXP3*, Tph-related double-positive cells exhibited reduced expression of *IL2RA* and *IKZF2*, alongside increased expression of *LAG3* and *KLRB1* (Figure 4D).

To validate these findings, we analyzed SF PD-1^+^HLA-DR^+^CD4^+^ T cells from o/p-JIA and sJIA patients by flow cytometry, stratifying cells based on granzyme A (GzmA) and FOXP3 expression. Among these activated T cells, the GzmA^−^FOXP3^−^subset, which is particularly enriched for Tph cells, was more prevalent in o/p-JIA, whereas the GzmA^−^FOXP3^+^ Treg population supposed to contain Tph-Treg double-positive cells was enriched in sJIA (Figure 4E). We then assessed CD25 (*IL2RA*) and CD161 (*KLRB1*) expression - markers differentially expressed in our transcriptomic analysis. The GzmA^−^FOXP3^+^ population in sJIA was skewed toward a CD161^+^CD25^−^ phenotype, aligning with the Tph-related double-positive subset identified by scRNA-seq (Figure 4E).

Collectively, these data support the expansion of a transcriptionally and phenotypically distinct population co-expressing Tph and Treg programs in sJIA, which may partially share differentiation pathways with Tph effector cells or even arise from Tph-like precursors.

### Dominant clonal expansion of effector over regulatory Tph cells in sJIA

Changes in CD4^+^ T cell subset frequencies provide only indirect insights into underlying biological processes such as local antigen encounter. Therefore, we next assessed clonal expansion within SF Tph cell subsets - either with an effector phenotype (Tph-only) or a potentially regulatory profile (double-positive) - as a more direct measure of local, antigen-driven activation.

TCR Vβ repertoire analysis revealed reduced clonal diversity among SF CD4^+^ T cells in sJIA compared to o/p-JIA, indicating more pronounced clonal expansion (Figure 5A and B). Stratification by cGEPs showed that the Tph^+^Treg^−^ double-positive subset in sJIA contained the highest frequency of expanded clones - exceeding all other CD4^+^ T cell subsets in both sJIA and o/p-JIA (Figure 5C). We next investigated the relationship between cGEP expression and clonal expansion. In sJIA, high Tph cGEP scores were closely associated with strongly expanded clones, while Treg-associated transcripts were primarily enriched in small to medium-sized clones and markedly reduced in highly expanded populations (Figure 5D). Cytotoxic and Th17 programs showed a more uniform distribution across clonal categories (Supplementary Figure 6). Across all clonal categories, Tph, Treg, and cytotoxic cGEP scores were elevated in sJIA relative to o/p-JIA, while Th17 scores were consistently lower in sJIA (Figure 5D, Supplementary Figure 6). Among clonally expanded cells, we could demonstrate clonal persistence over the disease course in two patients from whom serial SF samples were available. Clones longitudinally tracked across different time points exhibited higher Tph cGEP scores compared to those without signs of clonal persistence, while Treg cGEP intensity varied between patients and across time points (Supplementary Figure 7).

**Figure 5 –.**
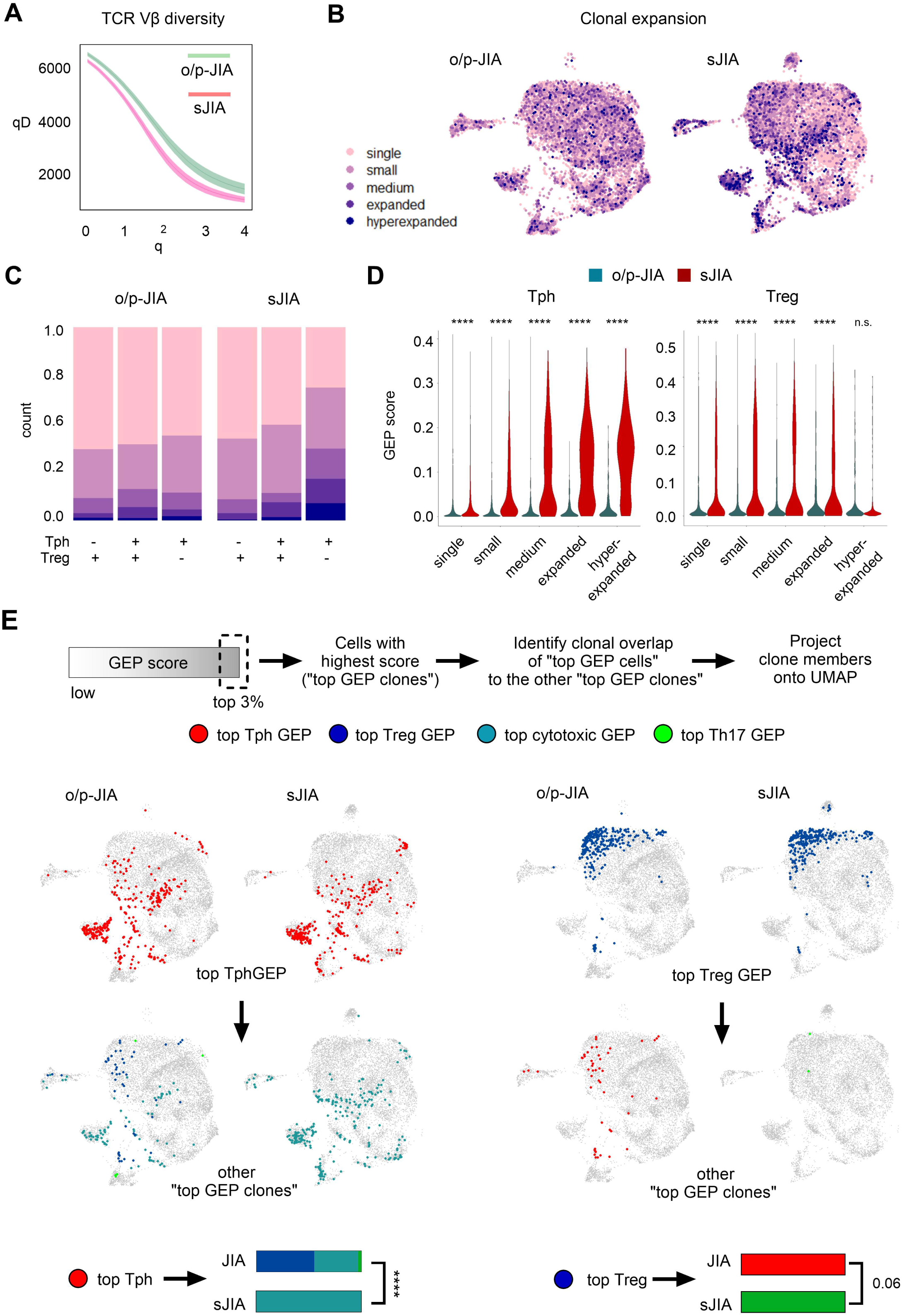
Clonal expansion favors effector over regulatory Tph cells in sJIA. (A) TCRβ repertoire diversity of SF CD4^+^ T cells from representative o/p-JIA and sJIA samples. Diversity index (qD) plotted across orders of diversity (q). (B) UMAP from Figure 2E annotated with clonal sizes. (C) Clonal distribution across subsets defined by Tph and/or Treg cGEP expression. (D) Correlation of clonal expansion with Tph and Treg cGEP scores. (E) Clonal relationships between highly polarized cells: top 3% of cells for each cGEP are mapped onto the UMAP, with shared clones in other polarized groups projected below. Comparisons between groups, as shown in the figure, were performed using Mann-Whitney test with Bonferroni adjustment for multiple testing in panel (D) and Fisher’s exact test in panel (E). p<0.001 (****), n.s., not significant.

Given the predominant convergence of Tph cGEP and clonal expansion in sJIA, we aimed at assessing the clonal relationship of highly polarized Tph cells to other functionally distinct subsets. For this, we identified the top 3% of cells with the highest expression of Tph, Treg, cytotoxic, or Th17 cGEP - representing the most transcriptionally polarized states. In sJIA, top-scoring Tph cells shared clonal overlap exclusively with cytotoxic top scorers, whereas in o/p-JIA, top-scoring Tph and Treg cells exhibited substantial clonal connectivity (Figure 5E). Notably, top-scoring Treg cells in sJIA lacked significant clonal overlap with any other subset, highlighting a clear divergence in lineage fate (Figure 5E).

These findings demonstrate that clonal expansion within the sJIA synovial CD4^+^ T cell compartment is driven predominantly by Tph^+^Treg^−^ effector cells. While in o/p-JIA, Tph and Treg cells may arise from shared precursors, potentially targeting the same antigen, sJIA is characterized by the selective clonal amplification of transcriptionally distinct Tph effector cells. This pattern supports a model of divergent differentiation in sJIA, favoring Tph-driven pathology in the inflamed joint.

### IL-1β and IL-18 promote Tph cell differentiation and cytokine secretion

Given the strong correlation between clonal expansion and a Tph-associated transcriptional signature in SF CD4^+^ T cells from sJIA patients, we investigated whether cytokines central to sJIA pathogenesis - specifically IL-1β and IL-18 - promote this effector program.

To test their role, we first stimulated CFSE-labeled naïve CD4^+^ T cells from healthy donors under limiting anti-CD28 co-stimulation, with or without IL-1β or IL-18. As expected, low CD28 signaling impaired proliferation, but both cytokines restored it under these suboptimal conditions, indicating a costimulatory function (Figure 6A, B).

**Figure 6 –.**
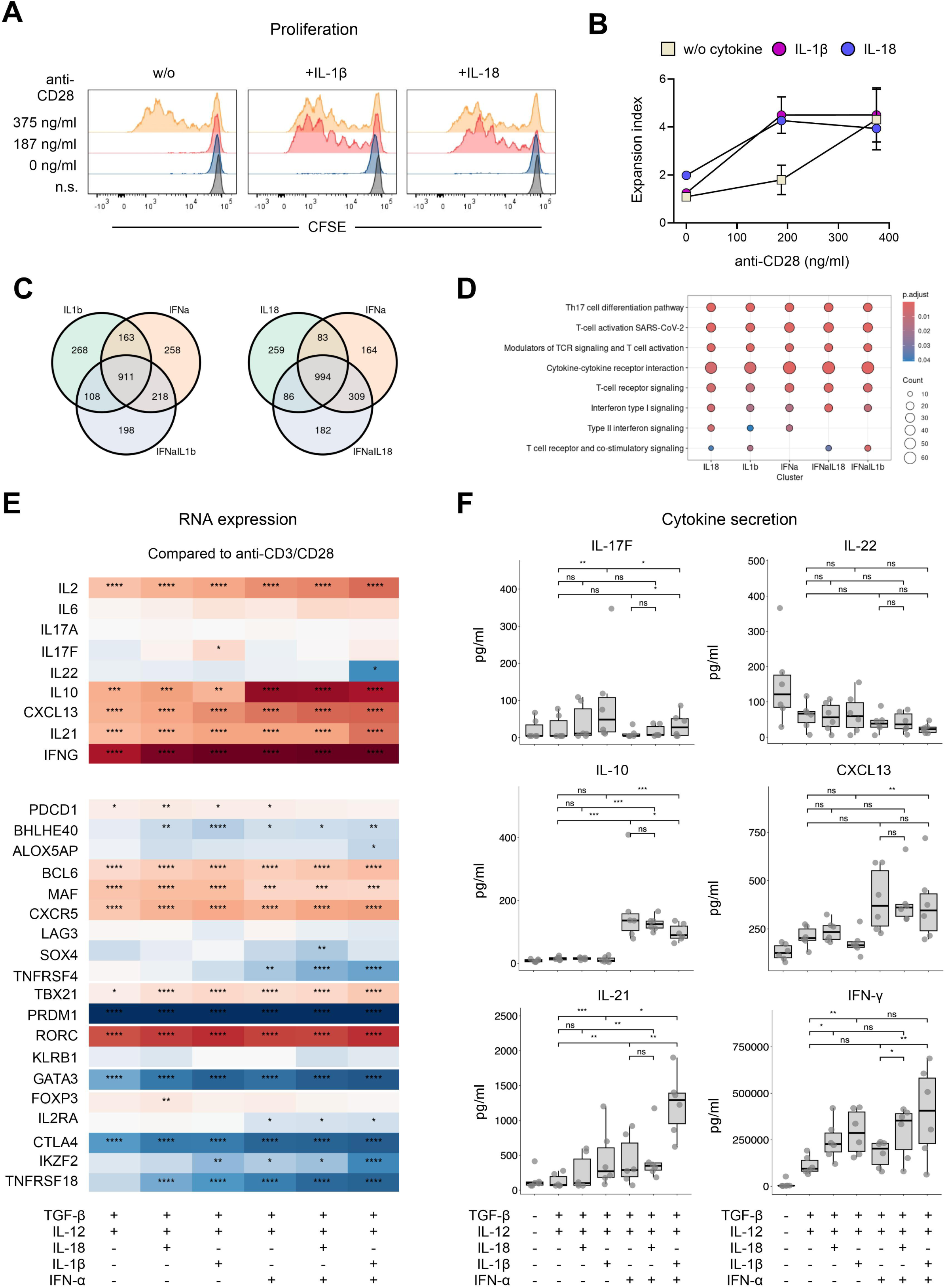
IL-1β and IL-18 provide co-stimulation and promote Tph cell differentiation and cytokine production. (A) Representative and (B) pooled data (n=3) showing proliferation of CFSE-labeled naïve CD4+ T cells from healthy individuals stimulated with anti-CD3 and titrated anti-CD28 ± IL-1β or IL-18. (C) Venn diagrams showing overlap of differentially expressed genes induced by IL-1β, IL-18, and IFN-α stimulation in naïve CD4+ T cells from healthy individuals (n=4) cultured under Tph-skewing conditions (TGF-β + IL-12). (E) Heatmaps showing gene expression profiles of naïve CD4+ T cells from healthy individuals (n=4) stimulated with IL-1β, IL-18 and/or IFN-α cultured under Tph-skewing conditions (TGF-β + IL-12), highlighting transcriptional signatures associated with T helper cell differentiation. Shown are Log_2_ fold changes compared to stimulation with anti-CD3/CD28 alone. (F) Cytokine concentrations in culture supernatants of naïve CD4+ T cells from healthy individuals (n=6) stimulated as described in (E). Comparisons between groups were performed as follows: two-tailed paired Student’s t-test in panel (B), Wald test implemented in DESeq2 in panel (E), and the Friedman test with Conover’s all-pairs comparison and Benjamini-Hochberg adjustment in panel (F). Data in (F) are presented as median with interquartile range and whiskers. Statistical significance is indicated as follows: p < 0.05 (*), p < 0.01 (**), p < 0.001 (***), p < 0.0001 (****); n.s., not significant.

To further dissect their role in Tph cell differentiation, we cultured naïve CD4^+^ T cells under Tph-skewing conditions (TGF-β + IL-12) and assessed the additional effects of IL-1β, IL-18, and IFN-α - either alone or in combination. IFN-α was included based on prior evidence of its capacity to enhance Tph polarization (26).

Transcriptomic analysis revealed substantial overlap in the differentially expressed gene (DEG) profiles induced by IL-1β, IL-18, and IFN-α, with a core gene set shared across all three conditions (Figure 6C). Pathway enrichment analysis showed that upregulated DEGs were predominantly linked to T cell receptor signaling and cytokine receptor activity. Of note, Th17 differentiation emerged as a consistently enriched pathway across all cytokine conditions (Figure 6D).

Hierarchical clustering of cytokine-induced transcripts (relative to anti-CD3/CD28 alone) identified gene modules synergistically upregulated by IL-1β or IL-18 in combination with IFN-α (Supplemental Figure 8 and 9). These clusters were enriched for canonical Tph-related genes, including *ICOS*, as well as effector cytokines *IL21* and *CXCL13* (Figure 6E; Supplementary Figure 8). In addition, Th1-associated transcripts such as *TBX21* and *IFNG* were co-enriched. Although upregulation of a subset of Th17-related genes was detected (*RORA*, *RORC, STAT3*), expression of hallmark Th17 cytokines were notably absent (Figure 6E). We next generated a cytokine-induced gene signature, capturing transcripts synergistically upregulated by IL-1β or IL-18 in combination with IFN-α, and mapped it onto the scRNA-seq data from SF CD4^+^ T cells in sJIA. This signature was preferentially enriched in Tph-only cells compared to both Treg-only and double-positive populations (Supplementary Figure 10). Since this effect is largely attributable to the expression of Tph effector cytokines such as CXCL13, the findings suggest that IL-1β and IL-18 predominantly drive cytokine effector programs in Tph cells lacking a regulatory profile.

To validate transcriptional findings at the protein level, we measured cytokine secretion in supernatants of the *in vitro* cultures. CXCL13 secretion was markedly enhanced in the presence of IFN-α, although neither IL-1β nor IL-18 further augmented this effect (Figure 6F). In contrast, IL-21 secretion was significantly enhanced by combined IFN-α and IL-1β stimulation, indicating a synergistic effect (Figure 6F). Consistent with transcriptomic data, IL-17 levels remained low.

Collectively, our data demonstrate that the sJIA-associated cytokines IL-1β and IL-18 enhance T cell proliferation and promote Tph cell effector molecules and cytokine secretion. These results establish a mechanistic link between the sJIA inflammatory milieu and the oligoclonal expansion of synovial Tph cells.

## Discussion

Our study reveals a marked expansion of activated CD4^+^ T cells in the SF of sJIA patients, reframing the role of adaptive immunity in a disease considered mainly driven by autoinflammatory mechanisms. Using flow cytometry, single-cell transcriptomics, clonal tracking, and *in vitro* differentiation assays, we show that antigen-experienced, clonally expanded CD4^+^ T cells in sJIA joints predominantly exhibit a Tph cell state. We further implicate IL-1β and IL-18 as key drivers of this cell state.

The expansion of Tph cells challenges the traditional dichotomy between autoinflammation and autoimmunity. Although sJIA is primarily driven by innate immune activation, particularly via IL-1β and IL-18, our data show that these cytokines also shape adaptive immunity by promoting the emergence of antigen-driven, pro-inflammatory CD4^+^ T cell subsets. This supports the biphasic model of sJIA, in which early IL-1-mediated inflammation modulates T cell responses, ultimately leading to autoimmune, T cell-driven arthritis (4). Our findings are consistent with a complementary study demonstrating that the complex SF cytokine matrix present in sJIA-affected joints preferentially drives T cell polarization towards Tph cells *in vitro* (27). In addition, earlier reports have shown that IL-1β directly enhances CD4^+^ T cell survival and effector function (28–30). Notably, IL-1β has been implicated in promoting T follicular helper (Tfh) cell activity within germinal centers (GCs) and augmenting IL-21 secretion via IL-1R1 signaling (31, 32).

In contrast to our findings, previous studies in sJIA have reported Th17-skewed signatures in CD4+ T cells, characterized by IL-17 expressing Tregs during early disease and a Th17-like transcriptional profile in effector T cells during later stages, despite limited IL-17 protein secretion (14). By comparison, our analysis reveals that clonally expanded CD4^+^ T cells in the joints are transcriptionally more aligned with Tph cells than with Th17 or other canonical T helper subsets.

Although these findings may seem contradictory at first glance, they may reflect overlapping biological programs. In our study, *in vitro* polarization under Tph-inducing conditions also led to the upregulation of some of these Th17-associated genes - including *RORA*, *RORC*, and *STAT3* - without eliciting IL-17 secretion. This suggests that certain effector T cell populations previously classified as Th17 based on transcriptional signatures may instead exhibit Tph-like characteristics. Consistent with this, naïve CD4^+^ T cells from sJIA patients readily differentiate into IL-21 producing Tfh-like cells, and motif enrichment analysis of blood samples identified the Tfh master regulator BCL6 as the most prominent transcription factor during active disease (18, 33). Alternatively, these differences may result from tissue compartmentalization, as peripheral blood T cell phenotypes may not accurately represent those present at inflammatory sites. However, across modalities - including scRNA-seq, flow cytometry, and functional assays - our data consistently point to a Tph-dominant phenotype in SF CD4^+^ T cells in sJIA, particularly among the clonally expanded populations indicative of local, antigen-driven activation.

The aforementioned observations raise the question of overlapping transcriptional features between Tph and Th17 programs, suggesting shared regulatory networks. Prior work demonstrated that TCR/CD28/Akt signaling strength in the presence of IL-1β influences T helper fate decisions, with strong signals suppressing Th17 differentiation while promoting Tfh-like expression profiles, including *ICOS* and *IL21* (*34, 35*). Moreover, in contexts of low or absent CD28 co-stimulation, IL-1β alone could support metabolic reprogramming necessary for T cell differentiation (34). Our *in vitro* data further show that IL-1β and IL-18 can compensate for limited CD28 signals to induce proliferation post-TCR engagement and, under full co-stimulation, in synergy with TGF-β, IL-12, and IFN-α enhance expression of Tph-associated genes, including *ICOS*, *IL21* and *CXCL13*.

Our data reveal a consistent enrichment of Tph cells in the inflamed joints of both o/p-JIA and sJIA. Despite this shared feature, subtle differences in the landscape of clonally expanded effector cells suggest divergent CD4^+^ T cell differentiation trajectories between the two disease forms. In sJIA, CD4^+^ T cells exhibit a stronger Tph-associated transcriptional signature. In contrast, a subset of effector cells in o/p-JIA expresses Th17-associated genes - an expression pattern largely absent in sJIA. These distinctions likely reflect the influence of distinct inflammatory milieus on T cell polarization and fate decisions (27, 36). The strength of integrated TCR/CD28 together with cytokine signaling appears to direct polarization toward Tfh or Th17 lineages (34). Strong co-stimulation, potentiated by IL-1β, favors the acquisition of Tfh-like features. Accordingly, elevated levels of IL-1β and IL-18 in sJIA joints may further reinforce local Tph differentiation (27, 36). IL-1β may also enhance the expression of costimulatory molecules on antigen-presenting cells, amplifying this effect (37). Notably, highly polarized Tph cells in sJIA were almost exclusively clonally linked to cytotoxic CD4^+^ T helper cells, a subset previously implicated in IL-18-driven responses (38). However, the precise contributions of IL-1β and IL-18 to the balance between these effector programs remain to be fully elucidated.

Strikingly, our analysis identified a distinct cell subset co-expressing Tph and Treg gene signatures, which was particularly expanded in sJIA. Similar Tph-like regulatory populations have previously been described in the SF of o/p-JIA patients and are thought to emerge from Tregs acquiring elements of the Tph transcriptional program (39, 40). These cells may resemble T follicular regulatory (Tfr) cells, known to restrain Tfh-driven B cell responses in GCs (41). IL-1β signaling is suggested to play a nuanced role in Tfh and Tfr cell biology, with differential expression of activating and inhibitory IL-1 receptors potentially allowing IL-1β to promote Tfh function at low concentrations and support Tfr cells at higher levels (31, 32, 42). Tfr cells comprise two subsets: natural Tfr cells, induced by IL-12 from thymus-derived Tregs and maintaining strong regulatory function; and induced Tfr (iTfr) cells, which arise from Tfh precursors and retain partial helper function (43, 44). In sJIA, a fraction of cells co-expressing Tph and Treg programs exhibited low *IL2RA* (CD25), alongside high *KLRB1* (CD161) and *LAG3* expression - features reminiscent of iTfr cells (44, 45). Their clonal relationship to Tph- only cells further supports a developmental trajectory along the Tph lineage. The marked expansion of Tph-related Tph^+^Treg^+^ cells in sJIA, together with the known impact of IL-1β on Tfh/Tfr dynamics, suggests these cells may represent a peripheral counterpart of iTfr cells, emerging in inflamed tissue under strong IL-1β–driven differentiation cues (32). However, their functional identity remains unresolved as we were unable to identify a distinct surface marker signature that would allow for specific isolation of this subset for functional analysis. This warrants further investigation to determine whether, in the chronically inflamed sJIA joint, their potential regulatory function is compromised, promoting the expansion of Tph-like effector cells.

Tph cells are key drivers of B cell activation across multiple autoimmune diseases (21, 22, 46, 47). In o/p-JIA, their expansion within the joints correlates with high ANA titers frequently observed already at disease onset (21, 39). Notably, both ANA-positive o/p-JIA and sJIA share a similar HLA background, particularly an increased prevalence of HLA-DRB1*11 alleles (11, 48). However, ANAs and other autoantibodies are typically absent at sJIA onset but may emerge later in the disease course (49, 50). These observations suggest that while Tph cells may contribute to autoantibody production in both diseases, the divergent timing of autoantibody development points to distinct mechanisms of B cell tolerance breakdown and highlights additional complexity in disease pathogenesis.

In summary, we reveal a dominant Tph transcriptional signature among clonally expanded CD4^+^ T cells in sJIA joints and demonstrate that IL-1β and IL-18 promote Tph differentiation and cytokine secretion. It suggests that sustained innate activation - particularly IL-1β and IL-18 - can act as a gateway for pathogenic T cell responses, establishing a mechanistic bridge between autoinflammation and autoimmunity. Therapeutically, this raises the possibility that blockade of these cytokines not only suppresses systemic inflammation but may also prevent the differentiation of pro-inflammatory Tph cells in the joints.

## Acknowledgements

We thank Gabriele Haase and Ursula Fischer for technical assistance. We thank the Core Unit for FACS and cell sorting of the IZKF Würzburg and the Single-Cell Center Würzburg for assistance with cell sorting and scRNA-seq.

## Contributors

JD, CK and HM designed the study. JD, JF and LH performed experiments. JD, JF, LH, CB, GP, MP, AHW, HJG, SMM, IC and CK collected data. JD, JF, LH, CB, HJG and CK analyzed data. JD and HM drafted the manuscript, and all authors revised and finally approved the manuscript.

## Ethics approval

The study was approved by the Research Ethics Committee of the University of Würzburg (299/17) as well as the Ethical Committee of Ospedale Pediatrico Bambino Gesù IRRCS in Rome (2333 OPBG 2020) and conducted in strict accordance with the principles of the Declaration of Helsinki. Written informed consent was obtained from the legal guardians of each participant.

## Data availability statement

The single cell and bulk RNA-sequencing data will be deposited in Gene Expression Omnibus and together with all other individual data are available from the corresponding author on reasonable request.

## Declaration of generative AI and AI-assisted technologies in the writing process

During the preparation of this work the authors used ChatGPT-4o in order to improve the readability and language of the manuscript. After using this tool, the authors reviewed and edited the content as needed and take full responsibility for the content of the publication.

## References

1. Pardeo M, Bracaglia C, De Benedetti F. Systemic juvenile idiopathic arthritis: New insights into pathogenesis and cytokine directed therapies. Best Pract Res Clin Rheumatol. 2017;31(4):505–16.

2. Erkens R, Esteban Y, Towe C, Schulert G, Vastert S. Pathogenesis and Treatment of Refractory Disease Courses in Systemic Juvenile Idiopathic Arthritis: Refractory Arthritis, Recurrent Macrophage Activation Syndrome and Chronic Lung Disease. Rheum Dis Clin North Am. 2021;47(4):585–606.

3. Kessel C, Hedrich CM, Foell D. Innately Adaptive or Truly Autoimmune: Is There Something Unique About Systemic Juvenile Idiopathic Arthritis? Arthritis Rheumatol. 2020;72(2):210–9.

4. Nigrovic PA. Review: is there a window of opportunity for treatment of systemic juvenile idiopathic arthritis? Arthritis Rheumatol. 2014;66(6):1405–13.

5. Erkens RGA, Calis JJA, Verwoerd A, De Roock S, Ter Haar NM, Den Engelsman G, et al. Recombinant Interleukin-1 Receptor Antagonist Is an Effective First-Line Treatment Strategy in New-Onset Systemic Juvenile Idiopathic Arthritis, Irrespective of HLA-DRB1 Background and IL1RN Variants. Arthritis Rheumatol. 2024;76(1):119–29.

6. Hinze CH, Foell D, Kessel C. Treatment of systemic juvenile idiopathic arthritis. Nat Rev Rheumatol. 2023;19(12):778–89.

7. Ter Haar NM, van Dijkhuizen EHP, Swart JF, van Royen-Kerkhof A, El Idrissi A, Leek AP, et al. Treatment to Target Using Recombinant Interleukin-1 Receptor Antagonist as First-Line Monotherapy in New-Onset Systemic Juvenile Idiopathic Arthritis: Results From a Five-Year Follow-Up Study. Arthritis Rheumatol. 2019;71(7):1163–73.

8. Pascual V, Allantaz F, Arce E, Punaro M, Banchereau J. Role of interleukin-1 (IL-1) in the pathogenesis of systemic onset juvenile idiopathic arthritis and clinical response to IL-1 blockade. J Exp Med. 2005;201(9):1479–86.

9. Quartier P, Allantaz F, Cimaz R, Pillet P, Messiaen C, Bardin C, et al. A multicentre, randomised, double-blind, placebo-controlled trial with the interleukin-1 receptor antagonist anakinra in patients with systemic-onset juvenile idiopathic arthritis (ANAJIS trial). Ann Rheum Dis. 2011;70(5):747–54.

10. Ombrello MJ, Arthur VL, Remmers EF, Hinks A, Tachmazidou I, Grom AA, et al. Genetic architecture distinguishes systemic juvenile idiopathic arthritis from other forms of juvenile idiopathic arthritis: clinical and therapeutic implications. Ann Rheum Dis. 2017;76(5):906–13.

11. Ombrello MJ, Remmers EF, Tachmazidou I, Grom A, Foell D, Haas JP, et al. HLA-DRB1*11 and variants of the MHC class II locus are strong risk factors for systemic juvenile idiopathic arthritis. Proc Natl Acad Sci U S A. 2015;112(52):15970–5.

12. Acosta-Rodriguez EV, Napolitani G, Lanzavecchia A, Sallusto F. Interleukins 1beta and 6 but not transforming growth factor-beta are essential for the differentiation of interleukin 17-producing human T helper cells. Nat Immunol. 2007;8(9):942–9.

13. Annunziato F, Cosmi L, Santarlasci V, Maggi L, Liotta F, Mazzinghi B, et al. Phenotypic and functional features of human Th17 cells. J Exp Med. 2007;204(8):1849–61.

14. Henderson LA, Hoyt KJ, Lee PY, Rao DA, Jonsson AH, Nguyen JP, et al. Th17 reprogramming of T cells in systemic juvenile idiopathic arthritis. JCI Insight. 2020;5(6).

15. Kessel C, Lippitz K, Weinhage T, Hinze C, Wittkowski H, Holzinger D, et al. Proinflammatory Cytokine Environments Can Drive Interleukin-17 Overexpression by gamma/delta T Cells in Systemic Juvenile Idiopathic Arthritis. Arthritis Rheumatol. 2017;69(7):1480–94.

16. Omoyinmi E, Hamaoui R, Pesenacker A, Nistala K, Moncrieffe H, Ursu S, et al. Th1 and Th17 cell subpopulations are enriched in the peripheral blood of patients with systemic juvenile idiopathic arthritis. Rheumatology (Oxford). 2012;51(10):1881–6.

17. Wilson NJ, Boniface K, Chan JR, McKenzie BS, Blumenschein WM, Mattson JD, et al. Development, cytokine profile and function of human interleukin 17-producing helper T cells. Nat Immunol. 2007;8(9):950–7.

18. Kuehn J, Schleifenbaum S, Hendling M, Siebenhandl S, Krainer J, Fuehner S, et al. Aberrant Naive CD4-Positive T Cell Differentiation in Systemic Juvenile Idiopathic Arthritis Committed to B Cell Help. Arthritis Rheumatol. 2023;75(5):826–41.

19. Yoshitomi H, Ueno H. Shared and distinct roles of T peripheral helper and T follicular helper cells in human diseases. Cell Mol Immunol. 2021;18(3):523–7.

20. Fischer J, Dirks J, Haase G, Holl-Wieden A, Hofmann C, Girschick H, et al. IL-21(+) CD4(+) T helper cells co-expressing IFN-gamma and TNF-alpha accumulate in the joints of antinuclear antibody positive patients with juvenile idiopathic arthritis. Clin Immunol. 2020;217:108484.

21. Fischer J, Dirks J, Klaussner J, Haase G, Holl-Wieden A, Hofmann C, et al. Effect of Clonally Expanded PD-1(high) CXCR5-CD4+ Peripheral T Helper Cells on B Cell Differentiation in the Joints of Patients With Antinuclear Antibody-Positive Juvenile Idiopathic Arthritis. Arthritis Rheumatol. 2022;74(1):150–62.

22. Rao DA, Gurish MF, Marshall JL, Slowikowski K, Fonseka CY, Liu Y, et al. Pathologically expanded peripheral T helper cell subset drives B cells in rheumatoid arthritis. Nature. 2017;542(7639):110–4.

23. Cano-Gamez E, Soskic B, Roumeliotis TI, So E, Smyth DJ, Baldrighi M, et al. Single-cell transcriptomics identifies an effectorness gradient shaping the response of CD4(+) T cells to cytokines. Nat Commun. 2020;11(1):1801.

24. Kotliar D, Curtis M, Agnew R, Weinand K, Nathan A, Baglaenko Y, et al. Reproducible single cell annotation of programs underlying T-cell subsets, activation states, and functions. bioRxiv. 2024.

25. Kotliar D, Veres A, Nagy MA, Tabrizi S, Hodis E, Melton DA, et al. Identifying gene expression programs of cell-type identity and cellular activity with single-cell RNA-Seq. Elife. 2019;8.

26. Tanemura S, Tsujimoto H, Seki N, Kojima S, Miyoshi F, Sugahara K, et al. Role of interferons (IFNs) in the differentiation of T peripheral helper (Tph) cells. Int Immunol. 2022;34(10):519–32.

27. Schleifenbaum, S, Swoboda, A., Dirks, J., Bracaglia, C., Hinze, T., Caiello, I., Prencipe, G., Pardeo, M., Park, C., Hinze, C., Wittkowski, H., Windschall, D., Foell, D., Morbach, H., Kessel, C. The systemic JIA synovial fluid environment supports development and prevalence of specific inflammatory T helper cell phenotypes. submitted. 2025.

28. Ben-Sasson SZ, Hu-Li J, Quiel J, Cauchetaux S, Ratner M, Shapira I, et al. IL-1 acts directly on CD4 T cells to enhance their antigen-driven expansion and differentiation. Proc Natl Acad Sci U S A. 2009;106(17):7119–24.

29. Jain A, Song R, Wakeland EK, Pasare C. T cell-intrinsic IL-1R signaling licenses effector cytokine production by memory CD4 T cells. Nat Commun. 2018;9(1):3185.

30. Park HJ, Shin MS, Shin JJ, Kim H, Kang B, Par-Young J, et al. IL-1 receptor 1 signaling shapes the development of viral antigen-specific CD4(+) T cell responses following COVID-19 mRNA vaccination. EBioMedicine. 2024;103:105114.

31. Belbezier A, Engeroff P, Fourcade G, Vantomme H, Vaineau R, Gouritin B, et al. Interleukin-1 regulates follicular T cells during the germinal center reaction. Front Immunol. 2024;15:1393096.

32. Vaineau R, Jeger-Madiot R, Ali-Moussa S, Prudhomme L, Debarnot H, Coatnoan N, et al. IL-1beta signaling modulates T follicular helper and regulatory cells in human lymphoid tissues. JCI Insight. 2025;10(12).

33. Hugle B, Schippers A, Fischer N, Ohl K, Denecke B, Ticconi F, et al. Transcription factor motif enrichment in whole transcriptome analysis identifies STAT4 and BCL6 as the most prominent binding motif in systemic juvenile idiopathic arthritis. Arthritis Res Ther. 2018;20(1):98.

34. Revu S, Wu J, Henkel M, Rittenhouse N, Menk A, Delgoffe GM, et al. IL-23 and IL-1beta Drive Human Th17 Cell Differentiation and Metabolic Reprogramming in Absence of CD28 Costimulation. Cell Rep. 2018;22(10):2642–53.

35. Bouguermouh S, Fortin G, Baba N, Rubio M, Sarfati M. CD28 co-stimulation down regulates Th17 development. PLoS One. 2009;4(3):e5087.

36. de Jager W, Hoppenreijs EP, Wulffraat NM, Wedderburn LR, Kuis W, Prakken BJ. Blood and synovial fluid cytokine signatures in patients with juvenile idiopathic arthritis: a cross-sectional study. Ann Rheum Dis. 2007;66(5):589–98.

37. Van Den Eeckhout B, Tavernier J, Gerlo S. Interleukin-1 as Innate Mediator of T Cell Immunity. Front Immunol. 2020;11:621931.

38. Barbosa CD, Canto FB, Gomes A, Brandao LM, Lima JR, Melo GA, et al. Cytotoxic CD4(+) T cells driven by T-cell intrinsic IL-18R/MyD88 signaling predominantly infiltrate Trypanosoma cruzi-infected hearts. Elife. 2022;11.

39. Jule AM, Lam KP, Taylor M, Hoyt KJ, Wei K, Gutierrez-Arcelus M, et al. Disordered T cell-B cell interactions in autoantibody-positive inflammatory arthritis. Front Immunol. 2022;13:1068399.

40. Lutter L, van der Wal MM, Brand EC, Maschmeyer P, Vastert S, Mashreghi MF, et al. Human regulatory T cells locally differentiate and are functionally heterogeneous within the inflamed arthritic joint. Clin Transl Immunology. 2022;11(10):e1420.

41. Fonseca VR, Ribeiro F, Graca L. T follicular regulatory (Tfr) cells: Dissecting the complexity of Tfr-cell compartments. Immunol Rev. 2019;288(1):112–27.

42. Pyrillou K, Humphry M, Kitt LA, Rodgers A, Nus M, Bennett MR, et al. Loss of T follicular regulatory cell-derived IL-1R2 augments germinal center reactions via increased IL-1. JCI Insight. 2024;9(5).

43. Castano D, Wang S, Atencio-Garcia S, Shields EJ, Rico MC, Sharpe H, et al. IL-12 drives the differentiation of human T follicular regulatory cells. Sci Immunol. 2024;9(97):eadf2047.

44. Le Coz C, Oldridge DA, Herati RS, De Luna N, Garifallou J, Cruz Cabrera E, et al. Human T follicular helper clones seed the germinal center-resident regulatory pool. Sci Immunol. 2023;8(82):eade8162.

45. Ritvo PG, Churlaud G, Quiniou V, Florez L, Brimaud F, Fourcade G, et al. T(fr) cells lack IL-2Ralpha but express decoy IL-1R2 and IL-1Ra and suppress the IL-1-dependent activation of T(fh) cells. Sci Immunol. 2017;2(15).

46. Law C, Wacleche VS, Cao Y, Pillai A, Sowerby J, Hancock B, et al. Interferon subverts an AHR-JUN axis to promote CXCL13(+) T cells in lupus. Nature. 2024;631(8022):857–66.

47. Dirks J, Fischer J, Klaussner J, Hofmann C, Holl-Wieden A, Buck V, et al. Disease-specific T cell receptors maintain pathogenic T helper cell responses in postinfectious Lyme arthritis. J Clin Invest. 2024;134(17).

48. Saleh R, Sundberg E, Olsson M, Tengvall K, Alfredsson L, Kockum I, et al. Genetic association of antinuclear antibodies with HLA in JIA patients: a Swedish cohort study. Pediatr Rheumatol Online J. 2024;22(1):79.

49. Hugle B, Hinze C, Lainka E, Fischer N, Haas JP. Development of positive antinuclear antibodies and rheumatoid factor in systemic juvenile idiopathic arthritis points toward an autoimmune phenotype later in the disease course. Pediatr Rheumatol Online J. 2014;12:28.

50. Krainer J, Hendling M, Siebenhandl S, Fuehner S, Kessel C, Verweyen E, et al. Patients with Systemic Juvenile Idiopathic Arthritis (SJIA) Show Differences in Autoantibody Signatures Based on Disease Activity. Biomolecules. 2023;13(9).

