## Supplementary Data for "Peripheral T Helper Cells Dominate the Synovial CD4^+^ T Cell Compartment in Systemic Juvenile Idiopathic Arthritis and Are Shaped by IL-1β and IL-18"

### SUPPLEMENTARY MATERIALS

### SUPPLEMENTAL METHODS

#### Patients and sample preparation

All patients were followed at the Division of Pediatric Rheumatology and Immunology, University Children's Hospital Würzburg, Würzburg, Germany or the Division of Rheumatology, Ospedale Pediatrico Bambino Gesù, IRCCS, Rome, Italy. Signed informed consent was obtained by the legal representatives. The study was reviewed by the Research Ethic Committee of the University of Würzburg (299/17) and the Ethical Committee of Ospedale Pediatrico Bambino Gesù IRRCS in Rome (2333 OPBG 2020) and conducted in accordance with the Declaration of Helsinki. Patients were not actively involved in the design, conduct, reporting, or dissemination plans of this research. Mononuclear cells were isolated from synovial fluid (SF) using Ficoll density-gradient centrifugation and stored in liquid nitrogen until use.

#### Antibodies, flow cytometry and cell sorting

Mononuclear cells were stained in 1X PBS 0.5% bovine serum albumin (BSA) with appropriate antibodies at 4°C for 30 minutes. Intracellular staining was carried out according to the manufacturer`s instruction using Intracellular Fixation & Permeabilization Buffer (eBioscience). For detection of intracellular cytokine expression, mononuclear cells were stimulated with phorbol 12-myristate 13-acetate (PMA, 50 ng/ml; Sigma-Aldrich) and ionomycin (1 µg/ml; Sigma-Aldrich) with addition of Brefeldin A (5 µg/ml, BioLegend) for 4 hours before staining. Flow cytometry data was acquired on a FACSCanto II (BD Biosciences) and analyzed with FlowJo version 10 (Tree Star).

Sorting of SF $CD4^+$ T cells and T cell subsets was performed on a FACSAria III (BD Biosciences). Alternatively, SF $CD4^+$ T cell have been purified using the EasySep Human CD4+ T Cell Isolation Kit (Stemcell Technologies). Peripheral blood (PB) B cells from healthy controls were immunomagnetically purified using CD20 microbeads (Miltenyi Biotec) and naïve CD4+ T cells using the EasySep Human Naïve T cell isolation Kit II (Stemcell Technologies).

The following antibodies were used for flow cytometry and cell sorting: CD3 BV510 (clone SK7), CD4 APC-Cy7 (clone OKT4), CD19 APC-Cy7 (clone HIB19), CD27 PerCP-Cy5.5 (clone M-T271), CD38 Brilliant Violet 421 (clone HIT2), CD25 APC (clone BC96), CD161 PerCP-Cy5.5 (clone HP-3G10), Granzyme A Pacific Blue (clone CB9), HLA-DR PE (clone L243), PD-1 PE-Cy7 (clone EH12.2H7), IL-21 APC (clone 3A3-N2), IFN-γ Brilliant Violet 510 (clone B27), TNF-α FITC (clone Mab11), IL-17 Brilliant Violet 421 (clone BL168) all from Biolegend and FOXP3 FITC (clone PCH101) from ebiosciences.

T cell/B cell co-culture assay

30,000 sorted SF $CD4^+$ PD-$1^+$ or PD-$1^-$ T cells were co-cultured with purified healthy control PB CD20+ B cells at a 1:1 ratio and stimulated with staphylococcal enterotoxin B (0.1 µg/ml) in round-bottom 96-well plates in 200µl RPMI 1640 medium supplemented with 10% FCS and penicillin/streptomycin. Differentiation of B cells into $CD19^+CD27^+CD38^{++}$ plasma cells was analyzed using flow cytometry after seven days.

T cell differentiation assay

Naïve $CD4^+$ T cells from healthy donors were resuspended in complete medium (RPMI 1640 with L-glutamine and sodium bicarbonate, 10% heat-inactivated FCS, penicillin-streptomycin, 10 mM HEPES, 50 µM b-mercaptoethanol, 1 mM sodium pyruvate, 10 µM non-essential amino acids) at a concentration of $1x10^6$/ml, transferred to a 96-well flat bottom suspension culture plate and stimulated with plate-bound anti-CD3 (clone HIT3a, Invitrogen) and anti-CD28 (clone CD28.2, BD Pharmingen) at a concentration of 3µg/ml each. Recombinant cytokines TGF-β1, IL-12p70, IFN-α 2a, IL-1β and/or IL-18 (all from Preprotech) were added, each at a final concentration of 10 ng/ml (equals 536 IU/ml for IFN-α 2a). Cells were cultured for 5 days at 37°C with 5% $CO_2$.

Bulk RNA-sequencing

RNA was isolated from four biological replicates of the T cell differentiation assay, each containing $2 \times 10^5$ T cells per condition, using the Qiagen RNeasy Micro Kit according to the manufacturer’s instructions. Sequencing was performed on the Illumina NovaSeq platform (Genewiz, Azenta Life Sciences).

Preprocessing of raw RNA sequencing data: After quality control and trimming with fastp version 0.23.4, raw reads were aligned to hg38 using STAR version 2.7.11a (1, 2). Sorting and indexing of aligned reads was performed using samtools version 1.18 and resulting read pairs were counted with featureCounts version 2.0.6 (3, 4).

Differential expression analysis: After filtering for protein coding genes above an expression threshold (10 read counts in at least 4 samples) DESeq2 version 1.42.1 in R version 4.3.2 was used for differential expression analysis. Log2 fold changes were calculated using Wald test with Benjamini-Hochberg adjustment (5). For visualization and ranking, log fold changes were shrunken using the ashr algorithm (6). Unless stated otherwise, a false discovery rate of 1 % (adjusted p value 0.01) was accepted for differential gene expression. We used the package clusterProfiler version 4.10.1 for discovery of enriched gene sets and pathways with a p value cut-off of 0.05 and q value cut-off of 0.2 (7).

ELISA

Cytokine concentrations were measured by ELISA for IL-21, CXCL13 and INF-γ using ELISA MAX Deluxe Set Human IL-21 (Biolegend), ELISA MAX Deluxe Set Human IFN-γ (Biolegend) and DuoSet ELISA Human CXCL13/BLC/BCA-1 (R&D Systems) and flow cytometry-based multiplex immunoassays using LEGENDplex HU Th17 Panel (7-plex) w/ FP V02 (Biolegend) for IL-17F, IL-22 and IL-10. For standard curve generation we used linear curve-fitting models for the single sandwich immunoassays, whereas we used the 5-parameter logistic regression fit (5PL) within the LEGENDplex Data Analysis Software, obtaining R2 values > 0.98 for all standard curves. Values below sensitivity limit or lower assay range limit were set to the respective limit value in sandwich immunoassays for CXCL13 and IL-21. For LEGENDplex data we set all values below the limit of detection, i.e. the minimum concentration that can be statistically distinguished from the background, to the respective value.

10X Genomics Chromium single-cell RNA-seq

Sorted SF CD4$^+$ T cells were subjected to scRNA-seq (10x Genomics). Samples were multiplexed with TotalSeqC Hashtag antibodies (Biolegend). After encapsulation and barcoding (10x Genomics), cells were lysed and cDNA prepared to create a 5′ gene expression library and a VDJ gene-enriched library for TCR repertoire analysis. Libraries were sequenced using Illumina NovaSeq 6000. Raw data were processed by CellRanger (10x Genomics) with standard settings. Subsequent analysis on scRNA-seq output files was carried out using Seurat version 5.0.1 in R version 4.3.2.

Preprocessing: Demultiplexing for JIA045 and JIA130 was done using cellsnp-lite and vireo and assigned to donors by combining this data with Hashtag Oligo UMIs in each donor. No multiplexing was used for sequencing of samples from JIA094 and JIA128. Remaining samples were demultiplexed using Hashtags and HTODemux within Seurat (positive.quantile = 0.99). Quality control was performed using manual thresholds. For bead-sorted samples (JIA094 and JIA128), RNA counts were required to range between 1,800 and 40,000, and detected features between 800 and 6,000. For all other FACS-sorted samples, RNA counts were set between 1,000 and 45,000, and features between 500 and 7,500. In all samples, mitochondrial read content was required to be below 15%. As an additional quality control measure, cells lacking a TCR sequence in the VDJ-seq data were excluded. Variable TCR genes were reassigned from the gene expression matrix to a separate assay sample, and the datasets were subsequently integrated using RPCA integration in Seurat following SCTransform normalization. A small cluster characterized by reduced CD4 expression, high expression of HLA-DR genes and *LYZ*, as well as elevated RNA and feature counts, was excluded as likely doublets with monocytes. The final data set included 10,216 and 9,777 cells from o-/p-JIA and sJIA patients, respectively.

Analysis: After reintegration clustering was done with FindClusters function of Seurat (resolution = .9). Differentially expressed genes (DEG) were identified by FindAllMarkers. Cluster annotation was performed using published scRNA-seq datasets of synovial $CD4^+$ T cells and compared to consensus gene expression programs (cGEPs) generated by the T-CellAnnoTator (TCAT) pipeline (8). The cGEP-scores per cell were integrated into the Seurat object for further analysis. TCR data was integrated from the CellRanger output airr tables. Expression of up to two different TCRα or β chains was accepted per cell. Multiple chains were ranked per UMIs. Clonotypes were defined as having the same V and J genes comprising the TCR and the same amino acid sequence of the CDR3 region. Clone sizes per sample were assigned as follows: 1 member: „single“, 2-5 members: „small“, 6-10 members: „medium“, 11-20 members „expanded“, > 20 members: „hyperexpanded”. For computation of diversity scores the alakazam package was used with standard settings. For covarying neighborhood analysis (CNA) the R implementation rcna of the python package cna was used (9). For generation of gene expression scores the AddModuleScore function of Seurat was used.

Data visualization

Figures related to scRNA-seq and bulk RNA-seq analysis were generated using various R packages, including alakazam, EnhancedVolcano, Nebulosa, ggplot2, ggalluvial, Seurat, and pheatmap. Quantitative data from other sources were visualized using Prism 10.1.0 (GraphPad).

Statistics

Statistical analyses were performed using Prism version 10.1.0 (GraphPad Software) and R (rstatix, dunn.test, ggpubr). For comparisons between two normally distributed groups, two-tailed unpaired or paired Student’s t-tests were used, or multiple t-tests with Bonferroni-Dunn correction for multiple comparisons where appropriate. For comparisons among more than two unpaired, normally distributed groups, one-way ANOVA followed by Bonferroni correction was applied. Results from these tests are presented as individual data points with mean ± standard deviation.For non-normally distributed data, the following non-parametric tests were employed: the Mann–Whitney U test with Bonferroni adjustment where appropriate for comparisons between two unpaired groups, and the Kruskal-Wallis test with Dunn correction for more than two unpaired groups. For more than two paired, non-normally distributed groups, the Friedman test was used followed by Conover’s all-pairs post hoc test with Benjamini–Hochberg adjustment. Data from non-parametric tests are presented as violin plots or median with interquartile range and whiskers. Fisher’s exact test was applied for analysis of categorical variables in contingency tables. A p-value < 0.05 was considered statistically significant.

**SUPPLEMENTARY TABLES**

| | **sJIA** (n= 7) | **o/p-JIA** (n= 22) | **ERA** (n=10) |
|---|---|---|---|
| **Age at onset** Median, interquartile range (years) | 5.8, 3.1 – 8.1 | 4.5, 2.5 – 9.0 | 14.1, 9.5 – 14.7 |
| **Age at sampling** Median, interquartile range (years) | 10.5, 6.2 – 11.4 | 9.7, 5.7 – 14.4 | 15.0, 14.4 – 15.8 |
| **Female** number (%) | 6 (75.0) | 18 (81.8) | 5 (50.0) |
| **ANA ≥ 1:160** number (%) | 1 (12.5) | 17 (77.3) | 4 (40.0) |
| **No treatment** number (%) | 0 (0.0) | 13 (59.1) | 5 (50.0) |
| **Only NSAID** number (%) | 0 (0.0) | 4 (18.2) | 4 (40.0) |
| **Glucocorticoids** number (%) | 4 (50.0) | 1 (4.5) | 0 (0.0) |
| **Sulfasalazine** number (%) | 0 (0.0) | 0 (0.0) | 1 (10.0) |
| **MTX** number (%) | 3 (37.5) | 4 (18.2) | 0 (0.0) |
| **TNF-α inhibition** number (%) | 1 (12.5) | 1 (4.5) | 0 (0.0) |
| **IL-1 inhibition** number (%) | 1 (12.5) | 0 (0.0) | 0 (0.0 |
| **IL-6 inhibition** number (%) | 4 (50.0) | 0 (0.0) | 0 (0.0) |

Supplementary Table 1 – Clinical and demographic characteristics of the study population

| Dg. | ID | sex | Age onset (years) | ANA | At sampling | | | |
|---|---|---|---|---|---|---|---|---|
| | | | | | **Age (years)** | **CRP (mg/dl)** | **ESR (mm/h)** | **Treatment** |
| sJIA | JIA151 | F | 5.9 | <1:80 | 10.3 | 5.8 | 42 | ETN, MTX, PDN |
| | | | | | 13.7 | 14.5 | 45 | ANK, MTX, PDN |
| sJIA | JIA157 | F | 5.5 | <1:80 | 10.1 | <0.05 | n.a. | TCZ |
| sJIA | JIA158 | M | 8.1 | <1:80 | 10.7 | 0.1 | 2 | TCZ |
| | | | | | 11.4 | <0.05 | 4 | TCZ |
| sJIA | JIA177 | M | 3.3 | <1:80 | 10.6 | <0.05 | 5 | TCZ, MTX, PDN |
| sJIA | JIA178 | F | 3.0 | <1:80 | 11.5 | 14.5 | 51 | MTX |
| p-JIA | JIA045 | F | 2.2 | 1:320 | 2.5 | <0.05 | 30 | NSAID |
| o-JIA | JIA094 | F | 4.3 | 1:320 | 4.8 | <0.05 | 11 | NSAID |
| o-JIA | JIA128 | F | 3.4 | 1:320 | 4.4 | <0.05 | n.a. | none |
| p-JIA | JIA130 | F | 1.5 | 1:160 | 7.5 | <0.05 | 6 | none |

Supplementary Table 2 – Clinical and demographical characterization of patients involved in scRNA-seq analylsis

Anakinra (ANK), Etanercept (ETN), Methotrexate (MTX), Prednisolone (PDN), Tocilizumab (TCZ), not assessed (n.a.)

### SUPPLEMENTARY FIGURES

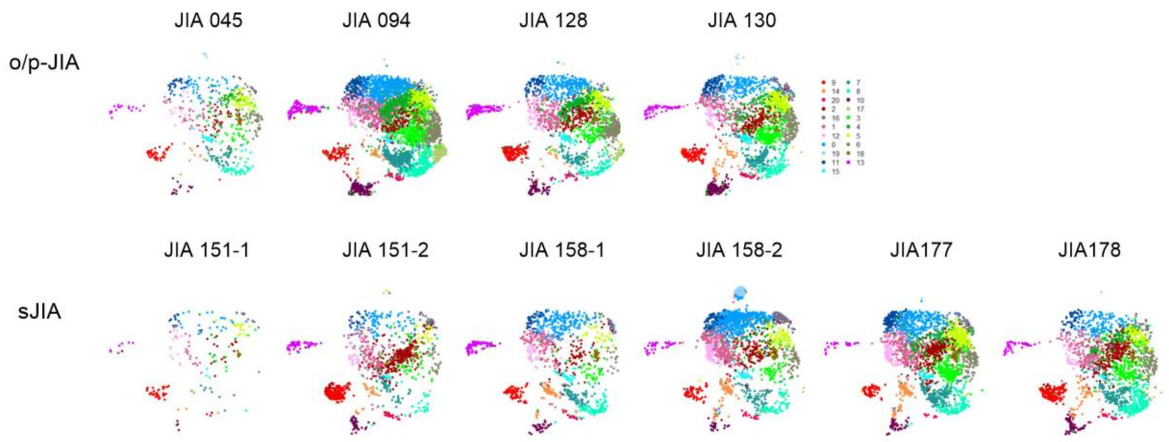


<u>Supplementary Figure 1 – Transcriptomic clusters are equally represented across patients</u>

UMAP projection of synovial fluid CD4$^{+}$ T cells with unsupervised cluster annotation, as in Figure 2A, displayed separately for each patient. Cells are colored by their assigned UMAP cluster. Synovial fluid samples from two sJIA patients were analyzed at two distinct time points during the disease course.

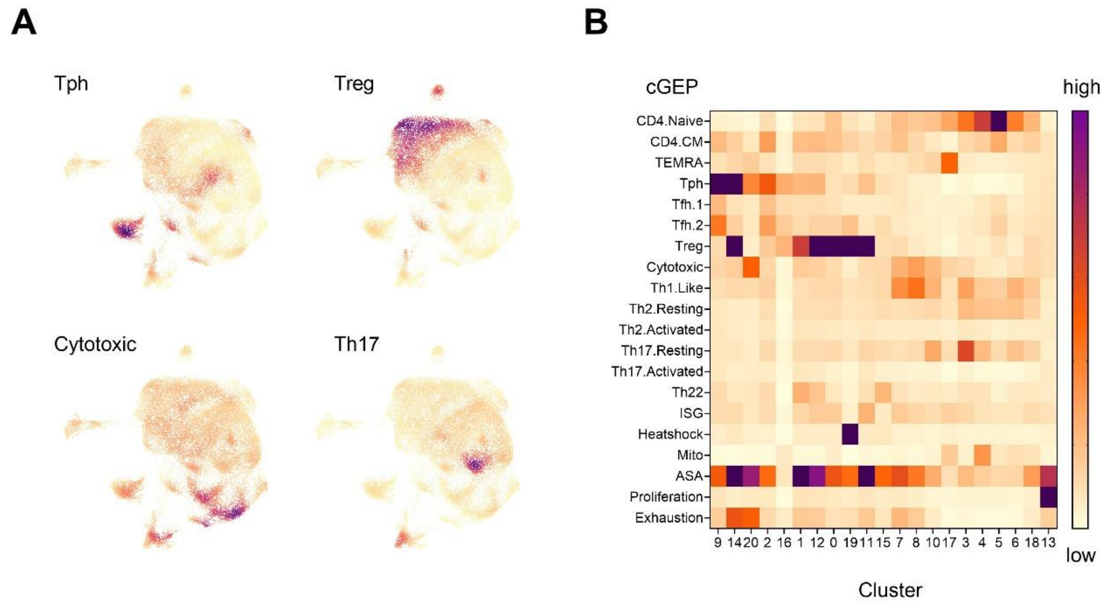


Supplementary Figure 2 – Overlapping usage of consensus gene expression programs across clusters

(A) UMAPs from Figure 2A, colored by selected consensus gene expression program (cGEP) usage. (B) Heatmap of cGEPs (rows) across clusters (columns).

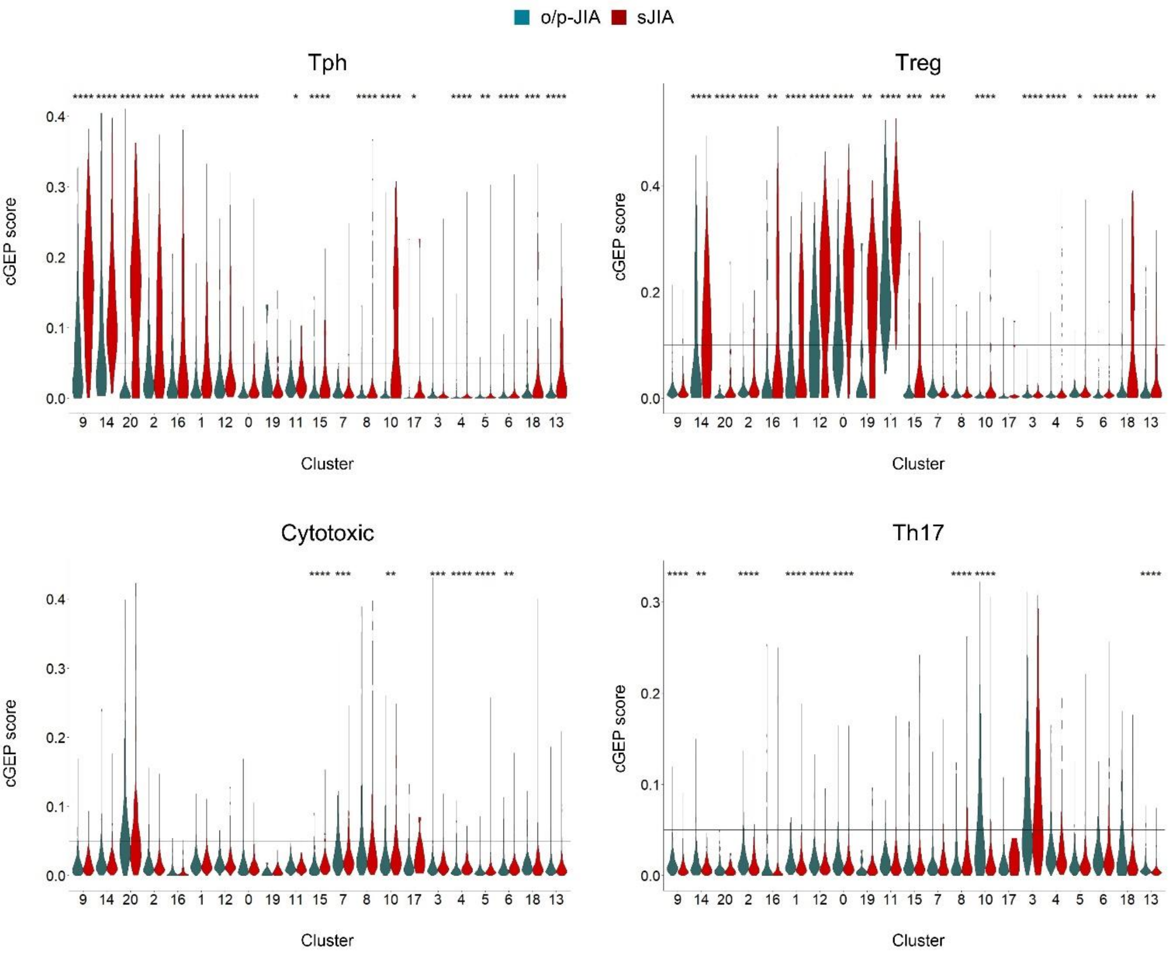


Supplementary Figure 3 – Dominant Tph gene expression programs partially overlap with Treg gene expression programs in sJIA

Violin plots depicting consensus gene expression programs (cGEPs) for each cluster of synovial fluid $CD4^+$ T cells of o/p-JIA (green) and sJIA (red) patients. Horizontal lines indicate the thresholds used to define cells as expressing the respective cGEP. Group comparisons, as shown in the figure, were performed using the Mann-Whitney test with Bonferroni correction for multiple comparisons. Statistical significance is indicated as follows: $p < 0.05$ (*), $p < 0.01$ (**), $p < 0.001$ (***), $p < 0.0001$ (****).

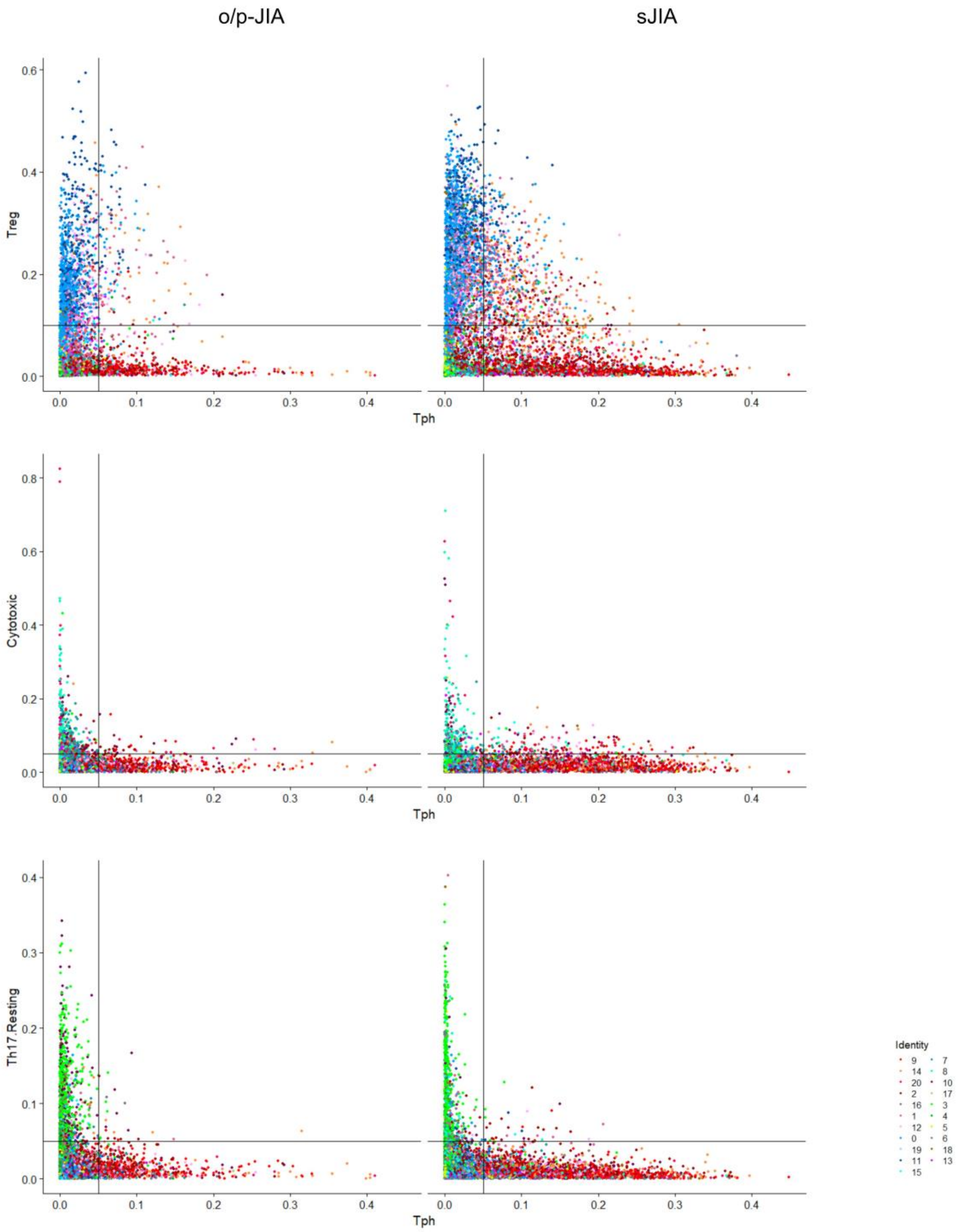


<u>Supplementary Figure 4 – Co-expression of Tph with other consensus gene expression programs</u>

Usage of the Tph against Treg, cytotoxic or Th17 consensus gene expression program (cGEP) in synovial fluid CD4$^+$ T cells from o/p-JIA or sJIA patients. Cells are colored by their assigned UMAP cluster, as defined in Figure 2A. Horizontal or vertical cut-offs indicate the thresholds for positivity for each respective cGEP.

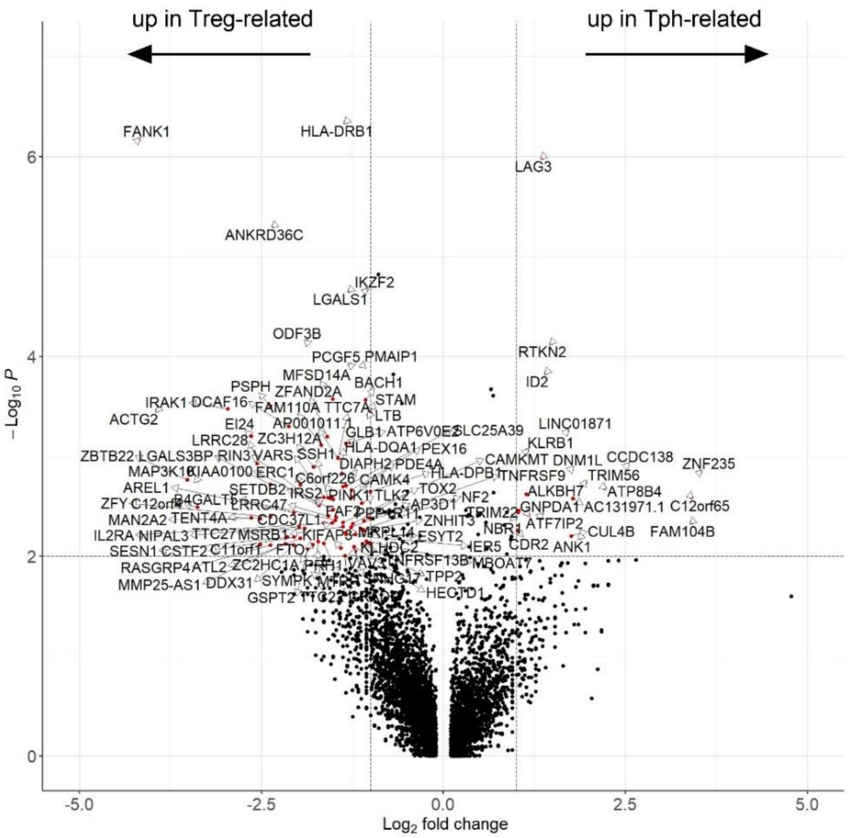


Supplementary Figure 5 - Differential gene expression between Tph-related and Treg-related Tph⁺Treg⁺ double-positive cells.

Volcano plot depicting differentially expressed genes between Tph- and Treg-related double-positive cells. Cells co-expressing both Tph and Treg consensus gene expression programs (cGEPs) were stratified based on clonal relationships: Tph-related double-positive cells share clones with cells expressing the Tph cGEP only, while Treg-related double-positive cells are clonally linked to cells expressing the Treg cGEP only. Genes with $Log_2$ fold change >1 and p-value <0.01 are highlighted in red.

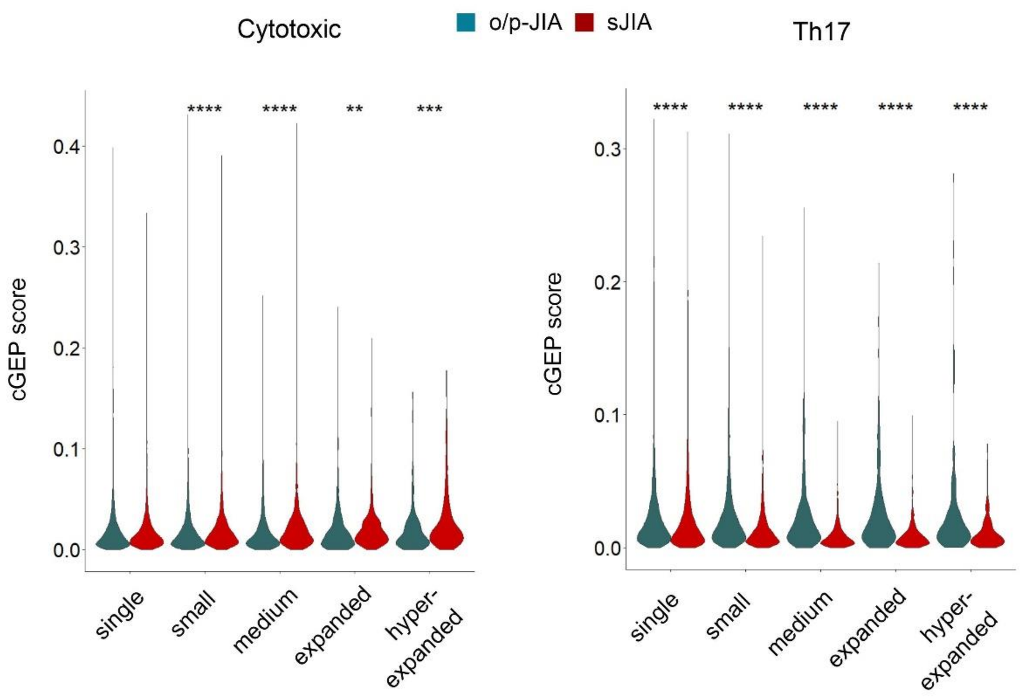


Supplementary Figure 6 – Correlation of cytotoxic and Th17 consensus gene expression programs with clonal expansion

Violin plots illustrating cytotoxic and Th17 consensus gene expression program (cGEP) scores in synovial fluid $CD4^{+}$ T cells, stratified by clonal size. Cells are grouped according to the extent of clonal expansion to assess the relationship between cGEP and clonal dynamics. Group comparisons, as shown in the figure, were performed using the Mann-Whitney test with Bonferroni correction for multiple comparisons. Statistical significance is indicated as follows: $p < 0.01$ (**), $p < 0.001$ (***), $p < 0.0001$ (****).

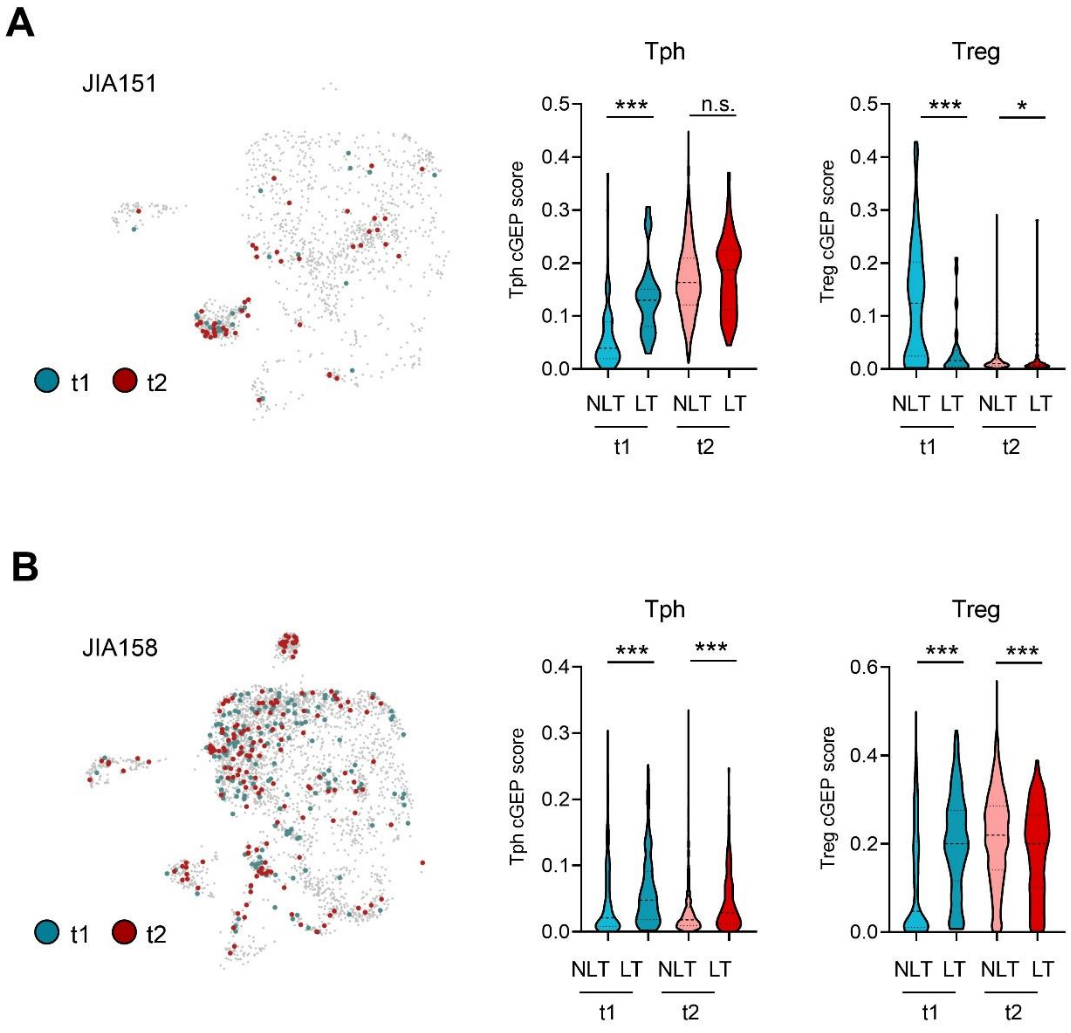


Supplementary Figure 7 – Tph and Treg gene expression programs during disease course

Consensus gene expression programs (cGEPs) for Tph and Treg cells were analyzed in clonally expanded $CD4^+$ T cells from synovial fluid samples of two sJIA patients, collected at distinct time points during the disease course. Clonally expanded cells (defined as ≥2 cells per clone within a patient) are color-coded by time point in the UMAP visualization. Within each patient and time point, cGEPs were compared between cells that were either clonally trackable across both time points (“longitudinally trackable”, LT) or not (“not longitudinally trackable”, NLT). Group comparisons were performed using the Mann–Whitney test. Statistical significance is indicated as follows: $p < 0.05$ (*), $p < 0.001$ (***); n.s., not significant.

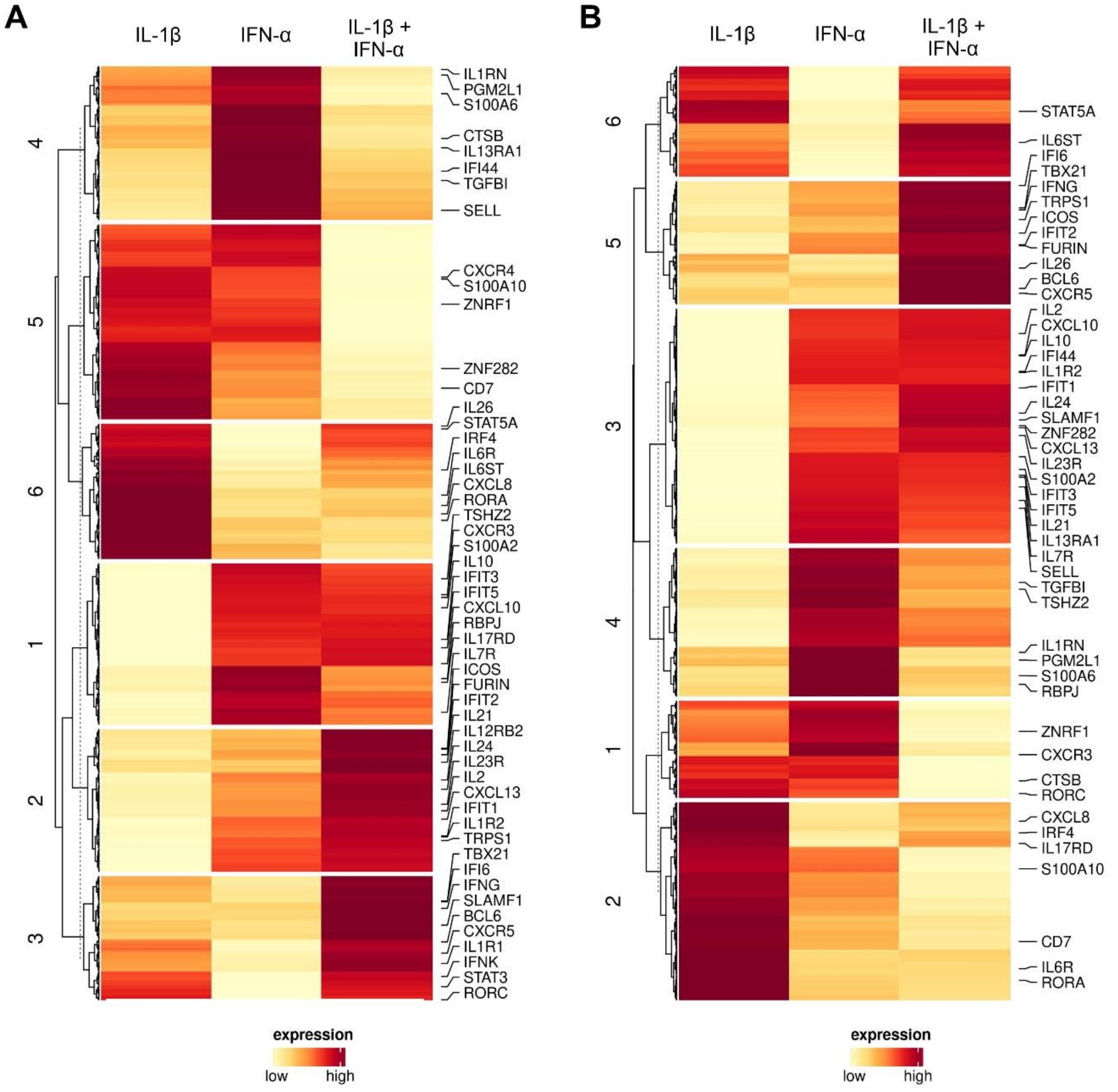


Supplementary Figure 8 – Upregulated transcriptional programs induced by IL-1β, IL-18, and IFN-α in TCR/CD28-activated naïve T cells

Naïve T cells were stimulated via anti-CD3/CD28 in the presence of different cytokine combinations for 5 days, followed by bulk RNA sequencing. The heatmap displays $Log_2$ fold changes in gene expression relative to stimulation with anti-CD3/CD28 alone. Only significantly upregulated genes (adjusted $p < 0.01$, $log_2FC > 0$) are shown. (A) Gene expression changes induced by IL-1β, IFN-α, or their combination. (B) Gene expression changes induced by IL-18, IFN-α, or their combination. Hierarchical clustering was performed using complete linkage and Euclidean distance metrics; dendrograms were cut into six clusters to define transcriptional modules.

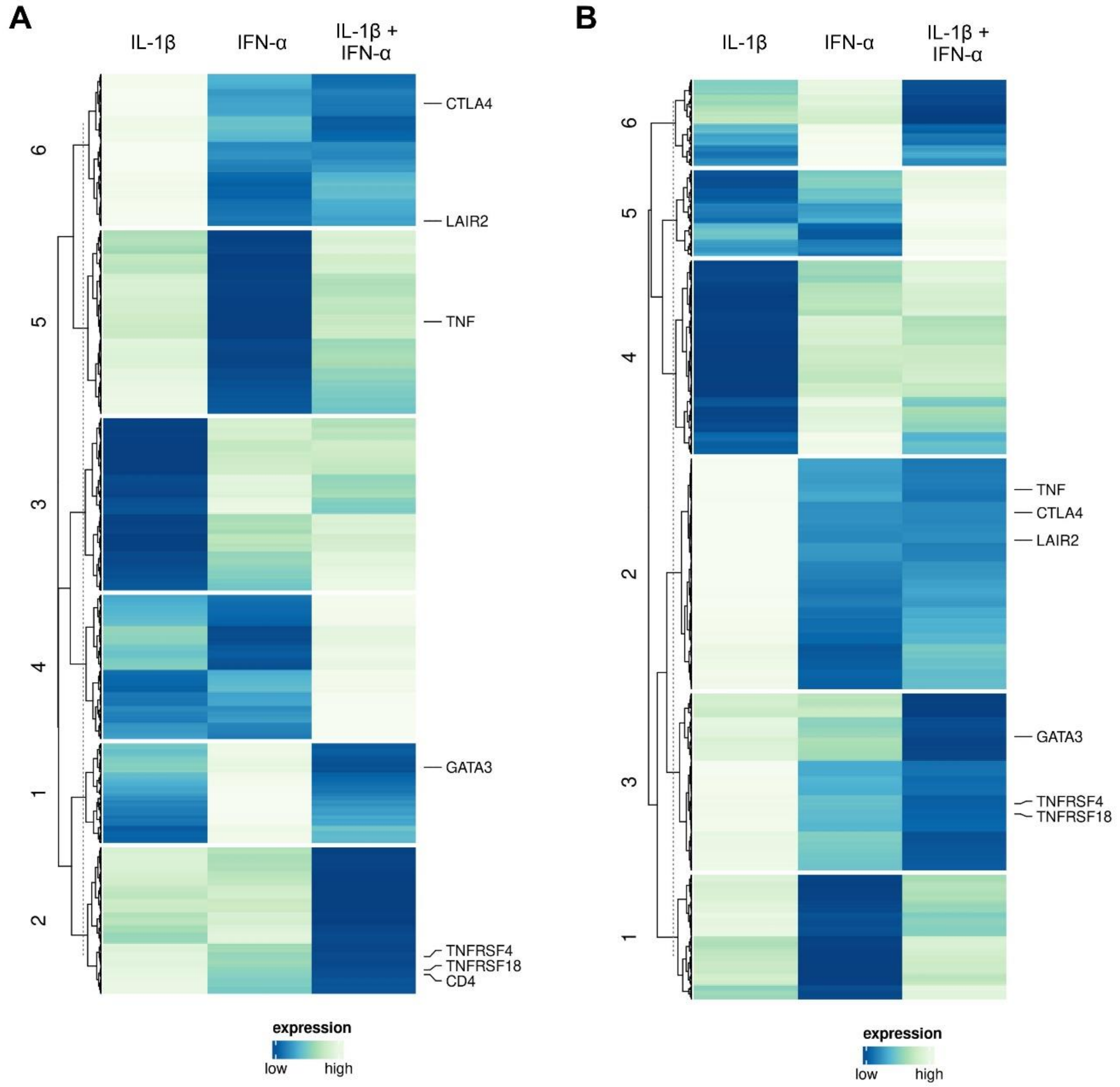


Supplementary Figure 9 – Downregulated transcriptional programs induced by IL-1β, IL-18, and IFN-α in TCR/CD28-activated naïve T cells

Naïve T cells were stimulated via anti-CD3/CD28 in the presence of different cytokine combinations for 5 days, followed by bulk RNA sequencing. The heatmap displays $\log_2$ fold changes in gene expression relative to stimulation with anti-CD3/CD28 alone. Only significantly downregulated genes (adjusted $p < 0.01$, $\log_2 FC > 0$) are shown. (A) Gene expression changes induced by IL-1β, IFN-α, or their combination. (B) Gene expression changes induced by IL-18, IFN-α, or their combination. Hierarchical clustering was performed using complete linkage and Euclidean distance metrics; dendrograms were cut into six clusters to define transcriptional modules.

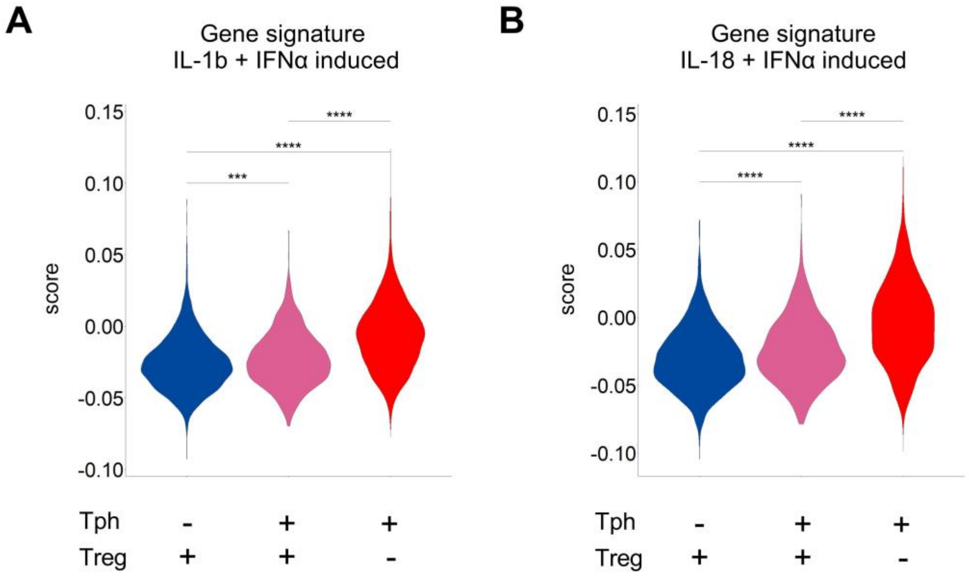


Supplementary Figure 10 – Expression of IL-1β/IFN-α and IL-18/IFN-α induced gene expression signatures mapped onto synovial fluid CD4$^{+}$ scRNA-seq data from sJIA patients

Violin plots comparing a gene signature derived from the synergistic effect of (A) IL-1β/IFN-α and (B) IL-18/IFN-α on antiCD3/CD28 stimulated naïve CD4+ T cells (data from Figure 6 and Supplementary Figure 8), with expression levels in synovial fluid CD4+ T cells from sJIA patients stratified by expression of a Tph and/or Treg consensus gene expression program (cGEP). Group comparisons, as shown in the figure, were performed using the Kruskal-Wallis test with Dunn correction for multiple comparisons. Statistical significance is indicated as follows: $p < 0.001$ (***), $p < 0.0001$ (****).

### Supplementary References


1. Chen S. Ultrafast one-pass FASTQ data preprocessing, quality control, and deduplication using fastp. Imeta. 2023;2(2):e107.
2. Dobin A, Davis CA, Schlesinger F, Drenkow J, Zaleski C, Jha S, et al. STAR: ultrafast universal RNA-seq aligner. Bioinformatics. 2013;29(1):15-21.
3. Danecek P, Bonfield JK, Liddle J, Marshall J, Ohan V, Pollard MO, et al. Twelve years of SAMtools and BCFtools. Gigascience. 2021;10(2).
4. Liao Y, Smyth GK, Shi W. featureCounts: an efficient general purpose program for assigning sequence reads to genomic features. Bioinformatics. 2014;30(7):923-30.
5. Love MI, Huber W, Anders S. Moderated estimation of fold change and dispersion for RNA-seq data with DESeq2. Genome Biol. 2014;15(12):550.
6. Stephens M. False discovery rates: a new deal. Biostatistics. 2017;18(2):275-94.
7. Wu T, Hu E, Xu S, Chen M, Guo P, Dai Z, et al. clusterProfiler 4.0: A universal enrichment tool for interpreting omics data. Innovation (Camb). 2021;2(3):100141.
8. Kotliar D, Curtis M, Agnew R, Weinand K, Nathan A, Baglaenko Y, et al. Reproducible single cell annotation of programs underlying T-cell subsets, activation states, and functions. bioRxiv. 2024.
9. Reshef YA, Rumker L, Kang JB, Nathan A, Korsunsky I, Asgari S, et al. Co-varying neighborhood analysis identifies cell populations associated with phenotypes of interest from single-cell transcriptomics. Nat Biotechnol. 2022;40(3):355-63.
